# The ciliary neurotrophic factor induces Stat3 phosphorylation in distinctive cytotypes of organs involved in body metabolism: an immunohistochemical study

**DOI:** 10.64898/2026.05.18.725839

**Authors:** Chiara Galli, Georgia Colleluori, Jessica Perugini, Edoardo Scopini, Ilenia Severi, Gaia Grandin, Antonio Giordano

**Affiliations:** Department of Experimental and Clinical Medicine, Marche Polytechnic University, 60126 Ancona, Italy; Center of Obesity, Marche Polytechnic University, 60126 Ancona, Italy; IRCCS INRCA, Ancona, Italy

**Keywords:** CNTF receptor, JAK-STAT, stomach, gut, pancreas, liver, adipose tissue, skeletal muscle

## Abstract

Administration of ciliary neurotrophic factor (CNTF) reduces food intake and body weight in both humans and experimental animals, where it also ameliorates hyperglycemia, hyperinsulinemia, and dyslipidemia. To exert its anti-obesogenic and anti-diabetogenic effects, CNTF targets brain feeding centers as well as multiple peripheral organs inducing the phosphorylation of the transcription factor signal transducer and activator of transcription 3 (p-STAT3). However, data showing which peripheral cytotypes are specifically targeted by exogenous CNTF *in vivo* in metabolically relevant organs are currently lacking. Here, we first evaluated the gene expression levels of the subunits of the tripartite CNTF receptor (Cntfr) complex, i.e., the *Cntfrα*, the leukemia inhibitory factor receptor β (*Lifrβ*) and the glycoprotein 130 (*gp130*), by quantitative real-time PCR in metabolically relevant organs of adult male mice: gastrointestinal (GI) tract, pancreas, liver, visceral and subcutaneous white (WAT) and interscapular brown adipose tissue (iBAT), skeletal muscle and the sciatic nerve. We then quantified p-STAT3 by Western blotting in these organs after intraperitoneal administration of CNTF (0.3 mg/kg) or saline. Finally, we mapped CNTF-responsive cells by immunohistochemistry, followed by morphometric quantification and confocal microscopy in both CNTF- and saline-treated mice. *Lifrβ* and *gp130* were ubiquitously detected across all the investigated organs; the *Cntfrα* showed the highest expression levels in the skeletal muscle, sciatic nerve, and iBAT, whereas it was found to be expressed to a lesser extent in the other sites. Administration of CNTF led to a significant increase of p-STAT3/STAT3 protein ratio in all organs examined, except the duodenum, and induced a distinctive pattern of cell nuclear p-STAT3 immunoreactivity. Notably, along the analyzed GI tract CNTF induced nuclear STAT3 phosphorylation in neurons of the submucosal and myenteric plexuses of the enteric nervous system and in contractile cells of the muscularis externa, where the response peaked in the mesenteric gut and colon. In the pancreas, CNTF triggered a higher activation within the endocrine component compared to the exocrine parenchyma. In the liver, CNTF induced STAT3 phosphorylation not only in parenchymal cells but also in sinusoids and resident macrophages. The cytokine activated p-STAT3 in subcutaneous and visceral white adipocytes, but also in brown adipocytes, with a prominent response observed in the beige subcutaneous adipocytes; adipose resident macrophages and endothelial cells of numerous blood vessels were also CNTF-responsive. Lastly, in skeletal muscle, a major site for glucose/lipid utilization, CNTF induced widespread nuclear p-STAT3 immunoreactivity in muscle fibers and in connective and Schwann cells of the peripheral nerves, including the sciatic nerve, supplying the gastrocnemius. In conclusion, our data indicate that CNTF acts across diverse cytotypes within metabolically relevant organs and tissues, likely fostering its peripheral metabolic effects through this cellular heterogeneity.

## 1. Introduction

The ciliary neurotrophic factor (CNTF) belongs to the interleukin (IL)-6 cytokine family, a group of structurally related cytokines including IL-6, IL-11, leukemia inhibitory factor (LIF), oncostatin M, cardiotrophin 1 (CT-1), cardiotrophin-like cytokine (CLC), and IL-27 (Rose-John, 2018). By autocrine, paracrine, and/or endocrine signaling, IL-6 cytokine family members affect several pathophysiological processes in mammals, ranging from the modulation of the immune response, especially B-cell stimulation and induction of the hepatic acute phase proteins, to neurotrophic effects and coordination of body metabolism (Miyaoka et al., 2006, Ghanemi and St-Amand, 2018, Murakami et al., 2019).

CNTF was originally discovered as a growth factor supporting the survival of chick ciliary ganglion neurons (Adler et al., 1979). Later, it was demonstrated to display significant neurotrophic effects for a wide variety of central and peripheral neurons, including motor neurons (Sendtner et al., 1994, Sleeman et al., 2000). To exploit possible therapeutic effects of CNTF, patients affected by amyotrophic lateral sclerosis were chronically treated with a recombinant human CNTF (Miller et al., 1996). These patients did not show amelioration of motor performance; however, they experienced anorexia and weight loss among other side effects. Subsequent clinical and pre-clinical studies confirmed that CNTF administration results in decreased food intake and weight loss and an improvement of obesity-associated hyperglycemia, hyperinsulinemia, and dyslipidemia in humans and in animal models (Lambert et al., 2001, Sleeman et al., 2003, Bluher et al., 2004). On a mechanistic level, administered CNTF reduces food intake by acting on hypothalamic and brainstem feeding centers (Lambert et al., 2001, Anderson et al., 2003, Senzacqua et al., 2016, Venema et al., 2020), but also improves obesity-associated metabolic alterations by acting on several metabolically relevant organs including liver, pancreas, skeletal muscle, and adipose tissue (Pasquin et al., 2015). However, a detailed anatomical description aimed at detecting which cytotypes are specifically targeted by exogenous CNTF in metabolically relevant organs is lacking.

The IL-6 family cytokines bind to multimeric receptor complexes (Rose-John, 2018). For CNTF, the cytokine binds to a three-part receptor complex consisting of the ligand-specific binding subunit receptor α (Cntfrα), the signal-transducing subunit (gp130), and the LIF receptor β (Lifrβ) (Davis et al., 1993, Ip et al., 1993). While gp130 is almost ubiquitously expressed by mammalian cells and the LIF receptor is common also to LIF, CT-1 and CLC signaling, the Cntfrα confers specificity to CNTF signaling (Sleeman et al., 2000). Importantly, CNTF activation of its tripartite receptor involves the activation of the janus family of tyrosine kinases (JAK1/JAK2) and signal transducers and activators of transcription (STAT), mainly STAT3 (Heinrich et al., 1998, Simi and Ibanez, 2010). STAT proteins are DNA-binding factors activated by JAK proteins through tyrosine phosphorylation and dimerization in response to extracellular signals, in turn causing the translocation of STAT dimers from the cytoplasm to the nucleus (Aaronson and Horvath, 2002). While systemic CNTF has a very short half-life (of the order of a few minutes as reported by (Dittrich et al., 1994)), the rapid nuclear translocation of p-STAT3, typically detected 45 minutes post-injection, provides a highly sensitive frame of the direct cellular response even after systemic clearance has occurred (Anderson et al., 2003; Kelly et al., 2004; Severi et al., 2012, 2013). Importantly, several studies previously demonstrated that STAT3 phosphorylation is induced by exogenous CNTF administration in both the central nervous system (CNS) (Severi et al., 2015, Venema et al., 2020) and in peripheral organs (Zvonic et al., 2003, Rezende et al., 2009, Perugini et al., 2019), and that this signaling pathway is the main responsible of CNTF effect (Rezende et al., 2009). The detection of the nuclear p-STAT3 immunoreactivity after a single CNTF injection is hence a reliable anatomical tool for the characterization of CNTF-responsive cells, similarly to what performed for the study of other cytokines including the transforming growth factor-β (Liu et al., 2013) and leptin (Hubschle et al., 2001, Frontini et al., 2008).

In the present study, CNTF was administered via a single intraperitoneal (i.p.) injection at a dose of 0.3 mg/kg, a paradigm widely established in the scientific literature (Lambert et al., 2001; Rezende et al., 2012) as it effectively mimics the anorectic and weight-regulatory effects of the cytokine without inducing the acute-phase inflammatory responses associated with higher dosages (Sleeman et al., 2003). With the primary aim of mapping cells responsive to exogenous CNTF at the anatomical level, we first quantified the expression of the tripartite receptor complex subunits *Cntfrα*, *Lifrβ* and *gp130* by quantitative real-time PCR (qRT-PCR) in murine peripheral organs playing crucial physiological functions in the regulation of the energy balance, such as nutrient digestion, absorption, metabolism, and storage (Yamada et al., 2008). We then quantified STAT3 activation by Western blotting and detected CNTF-responsive cells, *i.e.* cells bearing the functional CNTF receptor, by immunohistochemistry and morphometry in organs and tissues from CNTF-treated and saline-injected mice. Lastly, the phenotype of CNTF-responsive cells was assessed by double labelling and confocal microscopy using well established cell type markers. Collectively, our data characterized the specific cellular targets of CNTF within peripheral metabolic organs, providing a structural basis to understand its physiological role in the regulation of energy balance.

## 2. Materials & Methods

### 2.1 Animals and experimental conditions

Two-month-old male C57BL/6 mice were purchased from Charles River and housed individually under constant environmental conditions, with a 12-hour (h) light/dark cycle at 22°C and *ad libitum* access to standard chow diet and water. Animals were deeply anaesthetized with isoflurane and sacrificed in a fed state between 11:00 am and 12:00 pm. The principles of good laboratory animal care practice were followed, and experiments were conducted in accordance with the Council Directive 2010/63/EU of the European Parliament. All experiments were approved by the Italian Ministry of Health (authorization number: 96E38).

### 2.2 Tissue collection and processing

For qRT-PCR analysis, anesthetized animals (n=4) were decapitated. For Western blotting, mice received a single i.p. injection of either recombinant rat CNTF (0.3 mg/kg of body weight; R&D Systems, #557-NT; n=3) or an equal volume of saline (CTRL; n=3). The injected volumes of CNTF or saline were kept equal according to body weight and were performed using a Hamilton syringe. Forty-five minutes post-injection, treated and control mice were sacrificed by decapitation. For both experimental settings the following samples were collected: stomach, duodenum, mesenteric intestine (jejunum and ileum), colon, liver, pancreas, inguinal white adipose tissue (iWAT), epididymal white adipose tissue (eWAT), interscapular brown adipose tissue (iBAT), the gastrocnemius skeletal muscle, and the sciatic nerve. Following the dissection, all specimens were snap-frozen in liquid nitrogen and stored at −80°C until further use.

For morphological analyses, mice were treated with CNTF (n=3) or saline (CTRL; n=3) as described above. Forty-five minutes post-injection, animals were anesthetized and transcardially perfused with 4% paraformaldehyde in 0.1 M phosphate buffer (PB), pH 7.4 and the above listed organs were collected. After post-fixation in 4% paraformaldehyde for 24 h at 4°C, specimens were dehydrated with increasing alcohol concentration, cleared with xylol and embedded in paraffin.

### 2.3 qRT-PCR

Total RNA was extracted with Trizol reagent (Invitrogen; #15596018), purified, digested with ribonuclease-free deoxyribonuclease and concentrated using Total RNA purification kit (Norgen Biotek Corp.; #17250) according to the manufacturer’s instructions. For determination of mRNA levels, 1 μg of RNA was reverse-transcribed with the High-Capacity cDNA RT Kit with RNase Inhibitor (Applied BioSystems; #4374967) in a total volume of 20 μl. qRT-PCR was performed using TaqMan Gene Expression Assays and TaqMan Master Mix (Applied BioSystems; #4304437). All probes (Table S1) were purchased from Applied BioSystems. Reactions were carried out by Step One Plus Real Time PCR system (Applied BioSystems) using 50 ng cDNA in a final reaction volume of 10 μl. The thermal cycle protocol consisted of initial incubation at 95°C for 10 min followed by 40 cycles of 95°C for 15 s and 60°C for 20 s. All samples were run in duplicate. Samples not containing the cDNA template were included as negative controls in all experiments. TATA box-binding protein (Tbp) was selected as housekeeping gene to normalize gene expression. Relative mRNA expression was determined by the ΔCt method (2^-Δ^ ^Ct^).

### 2.4 Western blotting

Tissue lysates were prepared using a lysis buffer containing 50 mM Tris–HCl (pH 7.4), 1% NP-40, 1 mM EDTA, 150 mM NaCl, 1 mM sodium orthovanadate, 0.5% sodium deoxycholate, 0.1% SDS, 2 mM phenylmethylsulfonylfluoride, and 50 mg/ml aprotinin. Samples were centrifuged, and protein concentrations were determined by the Bradford Protein Assay (Bio-Rad Laboratories, Segrate, Italy). Equal amounts of protein were separated by SDS-PAGE and transferred onto nitrocellulose membranes using the Trans-Blot TurboTM Transfer system (Bio-Rad). To verify loading and transfer efficiency, membranes were visualized with Ponceau S solution (Santa Cruz Biotechnology, Santa Cruz, CA, USA). Membranes were then blocked for 1 h at room temperature (RT) in TBS-T (50 mM Tris-HCL [pH 7.6], 200 mM NaCl, and 0.1% Tween-20) containing 5% non-fat dried milk, followed by overnight incubation at 4°C with the primary antibody (Table S2a). After washing in TBS-T, membranes were incubated for 1 h at RT with the appropriate HRP-conjugated secondary antibody (Table S2b). Immunoreactive bands were visualized using the Clarity™ Western ECL substrate and the Chemidoc Imaging System (all from Bio-Rad). Densitometric analysis of bands was performed using Bio-Rad Image Lab software. Where appropriate, membranes were stripped, washed, and re-probed for total protein content.

### 2.5 Peroxidase immunohistochemistry

Peroxidase immunohistochemistry was performed on 4 µm thick paraffin-embedded sections. To minimize procedural variability in the detection of p-STAT3 immunoreactive cells, sections of organs from vehicle- and CNTF-treated mice were exposed to immunoperoxidase procedures in parallel. Sections were rehydrated and then subjected to an unmasking procedure at 95°C for 20 min in an antigen retrieval solution (Bio-Optica; #DV2004G1). Then, sections were treated with 3% H_2_O_2_ (in dH_2_O; 5 min) to block endogenous peroxidase, rinsed with phosphate-buffered saline (PBS), and incubated in a 3% normal goat serum (Vector Laboratories; #S-1000; in PBS, 20 min). Sections were then incubated overnight at 4°C with the monoclonal rabbit p-STAT3 antibody (Table S2a), which served as our primary reference for all peroxidase-based detections. After a thorough rinse in PBS, sections were incubated in the anti-rabbit IgG biotinylated solution (Table S2b) in PBS for 30 min. Histochemical reactions were performed using Vectastain ABC kit (ABC Kit Elite Peroxidase Standard, Vector Laboratories; #PK6100) and 3,3’-diaminobenzidine as substrate (DAB Substrate kit Peroxidase, Vector Laboratories; #SK4105). Sections were finally counterstained with haematoxylin, dehydrated, and mounted in Eukitt (Bio-Optica). To assess the specificity of the antibody, negative controls were obtained by omitting the primary antibody. The immunohistochemical staining was analysed using a Nikon Eclipse E600 microscope (Nikon).

### 2.6 Immunofluorescence and confocal microscopy

Immunofluorescence was performed on paraffin sections as follows: after dehydration, slices were washed in PBS containing 0.1% Tween for 5 min and then treated with a retrieval solution (Nacalai Tesque; #06380-05; 1:10 in dH_2_O; pH 7) at 70°C for 40 min. Then, sections were treated with a blocking solution (Nacalai Tesque; #06349-64) at RT for 40 min. Antibodies were applied overnight at 4°C. For immunofluorescence experiments, the choice between the monoclonal rabbit or the mouse p-STAT3 antibody (Table S2a) was determined by the host species of the specific cellular markers used for double or triple staining, ensuring the absence of cross-reactivity. Specifically, the mouse anti-p-STAT3 antibody was employed for co-localization studies in the mesenteric gut, liver, iWAT, and iBAT. The next day, sections were incubated in fluorophore-linked secondary antibodies solution (Table S2b) in PBS for 1h. Nuclear staining was performed using the fluorescent dye TO-PRO-3 Iodide (642 nm of excitation wavelength) (Invitrogen by Thermo Fisher; #T3602) in PBS for 15 min. Sections were subsequently mounted on standard glass slides and covered using Vectashield mounting medium (Vector Laboratories; #H-1000-10). Colocalization between antibodies was analysed by a motorized Leica DM6000 microscope at 40X and 60X magnifications. Fluorescence was detected with a Leica TCS-SL spectral confocal microscope equipped with an Argon and He/Ne mixed gas laser. Fluorophores were excited with the 488, 543 and 649 nm lasers and imaged separately. Images (1024 × 1024 pixels) were obtained sequentially from two channels using a confocal pinhole of 1.1200 and stored as TIFF files. The brightness and contrast of the final images were adjusted using Photoshop6 (Adobe Systems, RRID: SCR_014199).

### 2.7 Morphometric analysis

Morphometric analyses were performed on immunoperoxidase-stained sections counterstained with hematoxylin. The percentage of p-STAT3-positive nuclei on the total number of nuclei was calculated in the specimens from CNTF-treated (n=3) and CTRL (n=3) mice. For each organ, 3 to 5 non-consecutive sections (at least 50 μm apart) were randomly selected and imaged using a Nikon Eclipse E600 microscope at 40x or 60x magnification. Within each section, at least five randomly selected different areas were analyzed, and a total of 500 to 1000 nuclei for each organ were manually counted using ImageJ. To ensure unbiased quantification, all acquired images were assigned a unique numerical code and randomized by an independent researcher prior to analysis. Therefore, the investigator performed p-STAT3-positive cell counting blinded to the treatment groups until the final data decoding. Nuclei were classified as p-STAT3-positive based on the presence of a distinct, dark brown reaction product.

### 2.8 Statistical Analysis

Data are reported as mean ± standard error of the mean (SEM). Normality of data distribution was assessed using the Shapiro-Wilk test. For comparisons between two groups within a single tissue type, a two-tailed Student’s T-test was used. For comparisons involving multiple groups or organs/compartments, a One-Way ANOVA followed by Tukey’s post-hoc test was employed. Statistical significance was set at p < 0.05. All analyses were performed using GraphPad Prism 6 software (RRID: SCR_002798).

## 3. Results

### 3.1 Expression of *Cntfrα, Lifrβ,* and *gp130* transcripts in metabolically relevant peripheral organs

Among the metabolically relevant peripheral organs analyzed, the skeletal muscle displayed the highest expression of *Cntfrα*, followed by a substantial expression in the sciatic nerve and iBAT (Fig. 1a). Among the alimentary system organs, the liver and stomach showed the most prominent expression levels, whereas the duodenum, mesenteric gut, and colon displayed progressively lower transcript abundance (Fig. 1b). The quantification of pancreatic *Cntfrα* was hampered by the high intrinsic ribonuclease activity typical of this tissue (Azevedo-Pouly et al., 2014, Augereau et al., 2016, Al-Adsani et al., 2022), which precluded the recovery of high-quality mRNA. To circumvent this technical limitation, we performed a cross-check with the publicly available scRNA-seq dataset, *Tabula Muris* (Tabula Muris, 2018). This analysis confirmed a robust and selective expression of *Cntfrα* within the endocrine pancreatic islets, providing an independent validation of the receptor’s presence in this organ. Notably, these available datasets, documenting *Cntfrα* expression in the liver, skeletal muscle, and adipose tissues, were also consistent with our results. In this dataset, however, the lack of *Cntfrα* detection in the other organs (*i.e*., stomach) may be attributed to the relatively low expression levels of *Cntfrα* in the gastrointestinal tract (GI), as well as to the well-known lower sensitivity of scRNA-seq compared to qRT-PCR (Kolodziejczyk and Lonnberg, 2018). Collectively, these data highlight a pronounced different quantitative distribution of the *Cntfrα* transcript in metabolically relevant peripheral organs.

**Figure 1.**
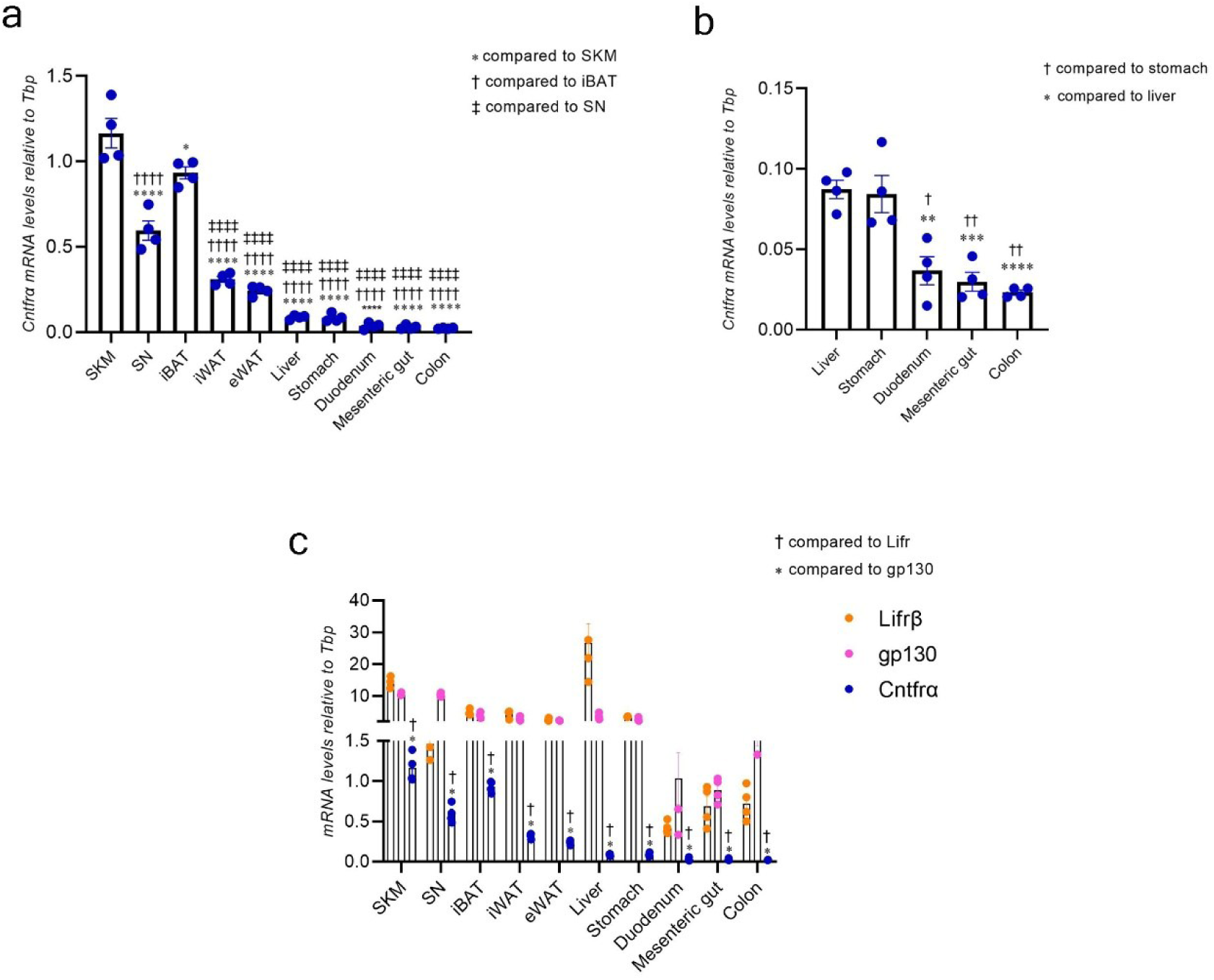
Cntfrα, Lifrβ and gp130 mRNA in murine metabolically relevant peripheral organs. (a) qRT-PCR analysis of *Cntfrα* expression levels in skeletal muscle (SKM), sciatic nerve (SN), interscapular brown adipose tissue (iBAT), inguinal white adipose tissue (iWAT), epididymal white adipose tissue (eWAT), liver, stomach, duodenum, mesenteric gut and colon. Panel (b) shows a higher magnification of the organs with the lowest expression levels of *Cntfrα* to better appreciate the differences between the liver, stomach, and intestinal segments (duodenum, mesenteric gut and colon). (c) Comparative analysis of the mRNA expression of *Cntfrα* (blue) with *Lifrβ* (orange) and *gp130* (magenta). Data are presented as mean ± SEM (n=4). In panels (a) and (b), the number of symbols indicates the level of significance: single p < 0.05; double p < 0.01; triple p < 0.001; quadruple p < 0.0001. In panel (c), statistical significance is indicated by a single symbol representing p < 0.0001 due to graph scale constraints. Data were analyzed using Two-way ANOVA followed by Tukey’s post-hoc test. *Tbp*=TATA-box binding protein.

Given that CNTF functional signaling requires the assembly of Cntfrα with the two signal-transducing subunits, gp130 and Lifrβ, we quantified their mRNA levels across all examined peripheral organs using qRT-PCR. Both *gp130* and *Lifrβ* displayed a quantitatively notable and ubiquitous expression throughout the peripheral tissues (Fig. 1c). This molecular profiling indicates that, while *Cntfrα* shows a more specific tissue distribution, the ubiquitous expression of *gp130* and *Lifrβ* identifies these organs as broad targets for the IL-6 family cytokines.

### 3.2 Detection and characterization of CNTF-responsive cells in metabolically relevant peripheral organs

To investigate the peripheral targets of exogenous CNTF, we mapped the cellular distribution of p-STAT3 across organs essential for nutrient handling, energy storage, and expenditure. Specifically, we performed a systematic anatomical and morphometric analysis of the GI tract, liver, pancreas, adipose tissues, and skeletal muscle. The anatomical description, combined with the results from our morphometric analyses, is summarized in Table 1.

**Table 1.** Summary of p-STAT3-positive nuclei distribution and quantification across peripheral organs.

| Organ | Anatomical Site | Cytotype | Marker | % of total p-STAT3(+) Nuclei - CTRL | % of total p-STAT3(+) Nuclei - CNTF | Δ % | p-value |
| --- | --- | --- | --- | --- | --- | --- | --- |
| Stomach | Submucosa | Connective and endothelial cells | Morphology | 6 ± 2% | 19 ± 3% | 13 ± 3% | p=0.0003 |
|  | Muscular layer | Smooth muscle cells | Morphology | 6 ± 2% | 29 ± 4% | 23 ± 5% | p<0.0001 |
| Duodenum | Submucosa | Brunner'sglands | Morphology | 6 ± 1% | 22 ± 3% | 16 ± 4% | p=0.0002 |
|  | Muscular layer | Smooth muscle cells | Morphology | 2 ± 1% | 38 ± 5% | 36 ± 7% | p=0.0001 |
| Mesenteric gut | Lamina propria | Connective, immune and endothelial cells | Morphology | 1 ± 0.4% | 27 ± 3% | 26 ± 4% | p<0.0001 |
|  | Submucosa | Connective and endothelial cells | Morphology | 5 ± 1% | 34 ± 2% | 30 ± 3% | p<0.0001 |
|  |  | Ganglia | PGP9.5 |  |  |  |  |
|  | Muscular layer | Smooth muscle cells | Morphology | 4 ± 1% | 47 ± 3% | 43 ± 4% | p<0.0001 |
|  |  | Ganglia | PGP9.5 |  |  |  |  |
| Colon | Lamina propria | Connective, immune and endothelial cells | Morphology | 0.2 ± 0% | 34 ± 6% | 34 ± 6% | p<0.0001 |
|  | Submucosa | Connective and endothelial cells | Morphology | 0.3 ± 0% | 30 ± 3% | 30 ± 3% | p<0.0001 |
|  |  | Ganglia | PGP9.5 |  |  |  |  |
|  | Muscular layer | Smooth muscle cells | Morphology | 1 ± 0% | 53 ± 5% | 52 ± 5% | p<0.0001 |
|  |  | Ganglia | PGP9.5 |  |  |  |  |
| Pancreas | Endocrine | α and β cells | Insulin / Glucagon | 2 ± 0% | 43 ± 5% | 41 ± 5% | p<0.0001 |
|  | Exocrine | Acinar cells | Morphology | 13 ± 3% | 26 ± 3% | 13 ± 5% | p=0.01 |
| Liver | Parenchyma | Hepatocytes | Morphology | 3 ± 1% | 53 ± 3% | 50 ± 4% | p<0.0001 |
|  | Sinusoids | Endothelial cells | Morphology |  |  |  |  |
|  |  | Kupffer cells | F4/80 |  |  |  |  |
| WAT | Inguinal | White adipocytes | PLIN-1 | 1 ± 0% | 44 ± 2% | 43 ± 2% | p<0.0001 |
|  |  | Macrophages | F4/80 |  |  |  |  |
|  |  | Endothelial cells | PECAM-1 |  |  |  |  |
|  |  | Beige adipocytes | Morphology | 0 ± 0% | 59 ± 2% | 59 ± 2% | p<0.0001 |
|  | Epididymal | White adipocytes | PLIN-1 | 3 ± 1% | 32 ± 2% | 29 ± 3% | p<0.0001 |
|  |  | Macrophages | F4/80 |  |  |  |  |
|  |  | Endothelial cells | PECAM-1 |  |  |  |  |
| BAT | Interscapular | Brown adipocytes | UCP1 | 0.4 ± 0% | 49 ± 2% | 49 ± 12% | p<0.0001 |
|  |  | Endothelial cells | Morphology |  |  |  |  |
| Skeletal muscle | Fibers | Myocytes | Morphology | 3 ± 1% | 36 ± 2% | 33 ± 3% | p<0.0001 |
|  | Sciatic nerve | Schwann's cells | S100b | 8 ± 1% | 45 ± 3% | 37 ± 4% | p<0.0001 |

Δ = (difference between means); (+) = positive; CTRL=saline-treated mice.

#### 3.1.1 Gastrointestinal tract

The CNTF administration triggered a widespread increase in STAT3 phosphorylation throughout the analyzed organs belonging to the GI tract.

In the stomach, Western blot analysis showed a very significant induction of p-STAT3 protein levels (Fig. 2a). Immunohistochemistry for p-STAT3 revealed a basal cytoplasmic immunoreactivity restricted to the gastric mucosa in the stomach of saline-treated mice (Fig. 2c), as previously reported (Judd et al., 2006). However, this basal signal remained unaffected by CNTF treatment (Fig. 2d). In contrast, CNTF specifically induced a significant p-STAT3 nuclear translocation within the smooth muscle cells of the muscularis externa, submucosal connective cells, and vascular endothelial cells (Fig. 2d). Quantitative morphometric analysis confirmed this activation, with p-STAT3-positive nuclei reaching 29 ± 4% (vs 6 ± 2% in the CTRL, p<0.0001) in the muscular layer and 19 ± 3% (vs 6 ± 2% in the CTRL, p=0.0003) in the submucosa (Fig. 2k and Table 1).

**Figure 2.**
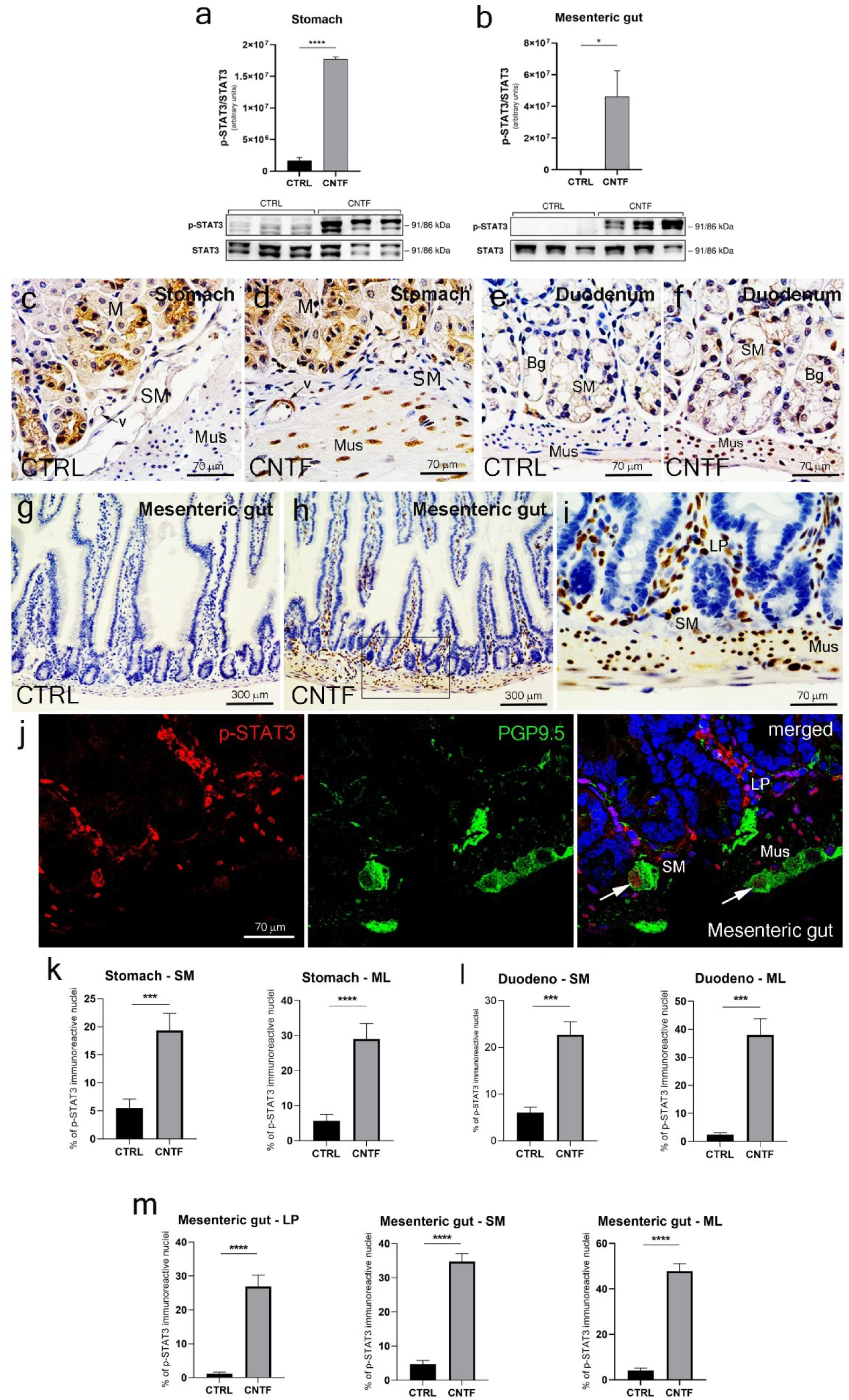
p-STAT3 immunoreactivity after CNTF administration in murine gastrointestinal tract. (a) Western blot analysis and corresponding densitometric quantification of p-STAT3 levels in the stomach and (b) mesenteric gut of saline- (CTRL) and CNTF-treated mice. Total STAT3 was used as a loading control. Peroxidase immunohistochemistry for p-STAT3 in the stomach (c-d), in the duodenum (e-f) and in the mesenteric gut (g-h) after saline or CNTF administration. Figure (i) is an enlargement of the corresponding areas framed in (h), showing a strong p-STAT3 immunoreactivity in several nuclei of the lamina propria and of the muscularis externa. Double-labelled confocal microscopy in the mesenteric gut of CNTF-treated mice (j), showing the colocalization between p-STAT3 and the pan-neural marker PGP9.5 in ganglia of the enteric nervous system in both the submucosal and myenteric plexuses (white arrows). Morphometric quantification of the percentage of p-STAT3-immunoreactive nuclei in the different anatomical layers of the stomach (k), duodenum (l), and mesenteric gut (m). Data are expressed as mean ± SEM (n=3 per group). *p < 0.05, ***p < 0.001, ****p < 0.0001 (Unpaired t-test). M: mucosa; SM: submucosa; Mus/ML: muscularis layer; LP: lamina propria; Bg: Brunner’s glands; v: blood vessels.

In the duodenum, Western blot analysis failed to show a significant increase of p-STAT3 protein levels; this lack of signal could not be attributed to protein degradation or poor sample quality, as the total STAT3 protein was clearly detectable and stable across all duodenal samples (data not shown). In striking contrast, while immunohistochemistry in saline-treated mice showed negligible p-STAT3 immunoreactivity, except for a faint staining in some submucosal Brunner’s gland cells (Fig. 2e), CNTF administration induced a widespread nuclear p-STAT3 immunoreactivity in cells of the muscularis externa (38 ± 5% vs 2 ± 1% in the CTRL, p=0.0001) and of the submucosal Brunner’s glands (22 ± 3% vs 6 ± 1% in the CTRL, p=0.0002; Fig. 2f, 2l and Table 1).

In the mesenteric gut, Western blotting revealed a significant increase of STAT3 phosphorylation following CNTF treatment (Fig. 2b). Consistently, our immunohistochemical assessment showed that CNTF induced a strong STAT3 phosphorylation in cells of the lamina propria (27 ± 2% vs 1 ± 0.4% in the CTRL, p<0.001; Fig. 2m and Table 1), the submucosa (34 ± 2% vs 0.2 ± 0% in the CTRL, p<0.001; Fig. 2m and Table 1), and the muscularis externa (47 ± 3% vs 4 ± 1% in the CTRL, p<0.001; Fig. 2m and Table 1). In these latter layers, p-STAT3 was also observed in PGP9.5-positive neurons of the submucosal and myenteric plexuses, respectively (Fig. 2j).

Similar results were obtained in the colon, where Western blotting revealed a marked increase in p-STAT3/STAT3 ratio upon CNTF treatment (Fig. S1a), which activated lamina propria, submucosa, and myenteric ganglia cells (53 ± 5% vs 1 ± 0% in the CTRL, p<0.0001; Figure S1b-d and Table 1). Both, the small and the large intestine contains a population of mucosal enteroendocrine cells, which are the main source of incretins, such as the glucagon-like peptide-1 (GLP-1) and gastric inhibitory peptide (GIP) (Hirasawa et al., 2005). GLP-1 and GIP are produced by epithelial cells of the mucosa, in particular GLP-1 is secreted by L-cells in both the small and the large intestine, while the GIP is mainly secreted by K-cells contained in the small intestine (Holst, 2004). However, we failed to observe p-STAT3 immunoreactivity in GLP-1- and GIP-positive cells in these organs after CNTF administration by double staining immunofluorescence and confocal microscopy (Fig. S2).

Overall, the present results indicate that administered CNTF does not act on epithelial cells lining the lumen of the analyzed GI tract segments. Instead, it targets specific cytotypes of the lamina propria, submucosa, muscularis externa, and the enteric nervous system, suggesting that CNTF influences gastrointestinal function indirectly by modulating, for example, Brunner’s gland cell physiology and gut motility rather than affecting epithelium-dependent secretory or absorbing functions.

#### 3.2.2 Pancreas

To evaluate the activation of the JAK/STAT3 pathway in the pancreas, we first performed Western blot analysis on total protein extracts. Systemic CNTF administration induced a significant increase in p-STAT3 protein levels compared to saline-treated mice, where the signal was markedly low (Fig. 3a). Immunohistochemistry in saline-treated mice revealed the presence of several p-STAT3-positive cells scattered in the exocrine compartment (data not shown), while the endocrine islets showed a nearly absent signal (Fig. 3b). Interestingly, morphometric analysis in saline-treated mice revealed a differential basal phosphorylation of STAT3 within the organ: the exocrine compartment displayed a significantly higher basal phosphorylation compared to the endocrine islets (13 ± 3% vs 2 ± 0%, respectively, p<0.05; Fig. 3e and Table 1). On the other hand, the magnitude of the response to systemic CNTF administration was markedly more evident in the pancreatic islets (compare Fig. 3c with 3b). In the endocrine compartment, CNTF induced a pronounced activation (reaching 43 ± 5% vs 2 ± 0% in the CTRL, p<0.0001; Fig. 3e and Table 1), whereas the response in the exocrine pancreas was comparatively lower, but still significant (26 ± 3% vs 13± 3% in the CTRL, p<0.05; Fig. 3e and Table 1). Murine pancreatic islets are composed primarily by insulin-secreting β-cells that in mice are clustered in a central core, surrounded by α-cells secreting glucagon, which is the second most represented cytotype in the pancreatic islets (Steiner et al., 2010). As shown by triple staining and confocal microscopy, CNTF induced p-STAT3 immunoreactivity in both insulin-positive β-cells and in glucagon-positive α-cells (Fig. 3d). In summary, our data indicate that endocrine islets represent the most responsive CNTF-target within the pancreas, while exocrine parenchymal cells display a weaker response.

**Figure 3.**
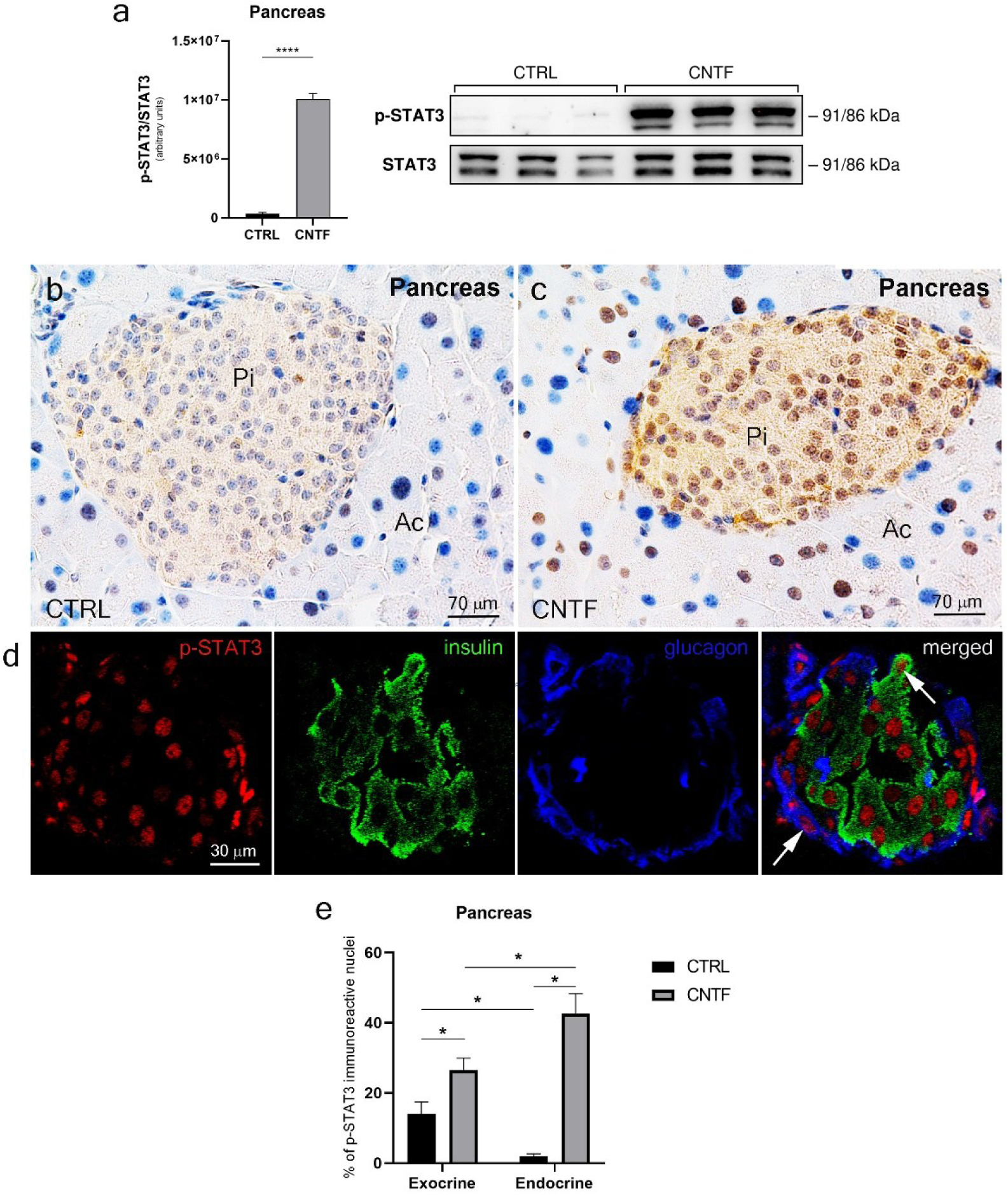
p-STAT3 immunoreactivity after CNTF administration in murine pancreas. (a) Representative Western blot and relative densitometric quantification of p-STAT3 protein levels in the pancreas of saline-(CTRL) and CNTF-treated mice. Total STAT3 was used as a loading control. (b-c) Peroxidase immunohistochemistry for p-STAT3 in pancreatic islets after saline or CNTF administration. (d) Double-labelled confocal microscopy for p-STAT3, insulin and glucagon (white arrows) in pancreatic islets after CNTF administration. (e) Morphometric quantification of p-STAT3-positive nuclei in the endocrine and exocrine components of the pancreas. Data are mean ± SEM (n=3 per group). *p<0.05; ****p < 0.0001; Unpaired t-test for (a); Two-way ANOVA followed by Tukey’s post-hoc test for (e). Pi= pancreatic islets; Ac= acinar cells.

#### 3.2.3 Liver

By Western blotting, in saline-treated mice p-STAT3 protein levels were virtually absent; CNTF administration induced a significant increase in STAT3 phosphorylation (Fig. 4a). Consistently, immunohistochemical analysis in control mice revealed only a weak cytoplasmic p-STAT3 immunoreactivity in the parenchyma of the liver, possibly due to non-specific background (Fig. 4b), while CNTF administration resulted in a strong STAT3 phosphorylation throughout the liver parenchyma (Fig. 4c). Such observations were confirmed by morphometric analysis revealing only very few p-STAT3 immunoreactive cell nuclei in saline-treated mice (Fig. 4e and Table 2), in stark contrast with a massive and widespread response in CNTF-treated mice (53 ± 3% vs 3 ± 1% in the CTRL, p < 0.0001; Fig. 4e and Table 1). The p-STAT3 immunoreactivity was predominantly localized in central and rounded nuclei of numerous polyhedral hepatocytes, forming the typical plates spaced by sinusoids (Fig. 4c). Binucleated hepatocytes and hepatocytes containing large, likely polyploid nuclei were also p-STAT3-positive (Fig. 4c, inset). In CNTF-treated mice, p-STAT3 immunoreactivity was also detected in other cell types than hepatocytes, including cells lining the wall of the sinusoids (endothelial cells) and other cells attached to the wall of the sinusoids (Fig. 4c, inset). Based on their shape and location, these latter CNTF-responsive cells resembled the resident macrophage population, also known as Kupffer’s cells (Naito et al., 2004). CNTF response in Kupffer’s cells was indeed confirmed by the presence of p-STAT3 immunoreactive nuclei in some cells positive for the macrophage marker F4/80 (Lumeng et al., 2007) (Fig. 4d). In conclusion, our data demonstrate that different cytotypes within the liver are responsive to systemic CNTF, given the profound induction of STAT3 phosphorylation in both the hepatocyte population and the sinusoidal immune-vascular component.

**Figure 4.**
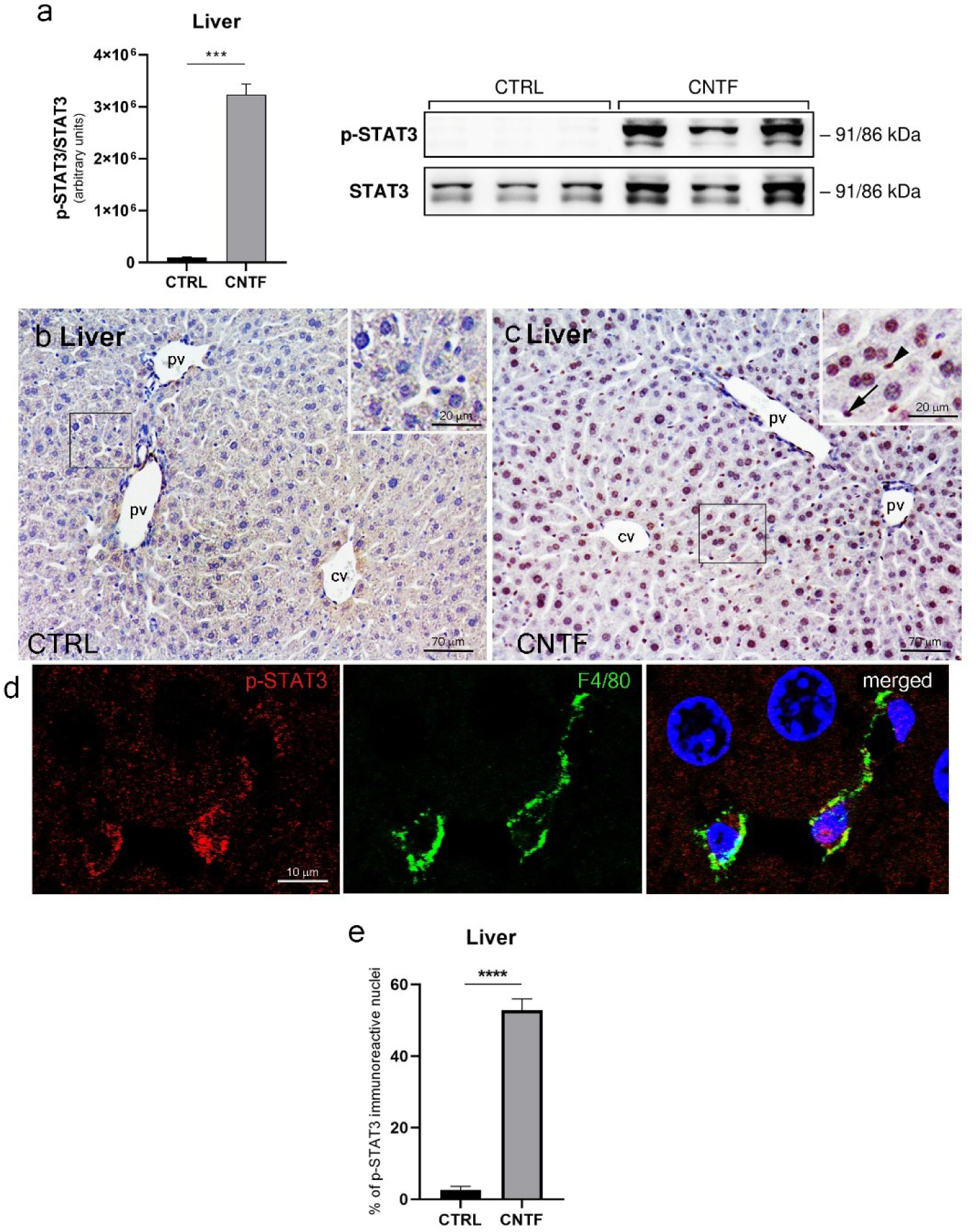
p-STAT3 immunoreactivity after CNTF administration in murine liver. (a) Representative Western blot and relative densitometric quantification of p-STAT3 protein levels in the liver of saline- (CTRL) and CNTF-treated mice. Total STAT3 was used as a loading control. (b-c) Peroxidase immunohistochemistry for p-STAT3 in the liver after saline or CNTF administration. Insets are enlargements of the corresponding framed areas in (b) and in (c). Figure (c) shows p-STAT3 immunoreactivity induced by CNTF in polyploid and binucleated hepatocytes, in elongated cells at the wall of the hepatic sinusoids (arrowhead), compatible with endothelial cells, and in group of cells located inside the hepatic sinusoids (black arrow), resembling Kupffer’s cells. (d) Double-labelled confocal microscopy for p-STAT3 and the macrophage-marker F4/80 in the liver after CNTF administration. (e) Morphometric quantification of p-STAT3-positive nuclei in the liver. Data are mean ± SEM (n=3 per group). ***p < 0.001; ****p <0,0001; Unpaired t-test; cv=central vein; pv=portal vein.

#### 3.2.4 Adipose tissues

The epididymal fat depot surrounding the rodent male reproductive organs was collected as representative of the visceral WAT. Visceral WAT is the main site of lipid storage in mammals and its expansion under conditions of increased caloric intake and/or reduced energy expenditure is strongly linked to the pathophysiology of obesity (Lumeng et al., 2007). Visceral WAT expansion progressively promotes liver metabolic dysfunction, systemic inflammation and insulin resistance, thereby significantly contributing to morbid obesity and multi-organ associated diseases (Lumeng et al., 2007). To evaluate the effect of the CNTF on this depot, we first analyzed eWAT total protein extracts *via* Western blotting. In saline-treated mice p-STAT3 was nearly undetectable, whereas CNTF administration induced a significant increase in protein phosphorylation (Fig. 5a). This activation was further detailed through immunohistochemical analysis: after saline treatment, p-STAT3-positive nuclei were not detected (Fig. 5b), while CNTF administration resulted in a strong p-STAT3 activation in eWAT (Fig. 5c). Accordingly, morphometric analysis revealed a nearly absent activation of p-STAT3 in saline-treated mice (Fig. 5g and Table 1), different from the widespread activation in CNTF-treated mice, where the percentage of p-STAT3-reactive nuclei rose significantly to 32 ± 2% (vs 3 ± 1% in the CTRL, p<0.0001; Fig. 5g and Table 1). Most of the CNTF-responsive cells were white adipocytes, as demonstrated by the presence of p-STAT3 immunoreactive nuclei at the periphery of large unilocular cells, that were also positive for the lipid droplet-associated protein perilipin-1 (PLIN-1) (Fig. 5d). p-STAT3 immunoreactivity was observed also in the majority of cells positive for the macrophage marker F4/80 (Fig. 5e) and in endothelial cells lining the lumen of numerous small blood vessels (arterioles, venules, and capillaries) and positive for the Platelet/Endothelial Cell Adhesion Molecule-1 (PECAM-1), an antigen expressed at the surface of endothelial cells (Lertkiatmongkol et al., 2016) (Fig. 5f).

**Figure 5.**
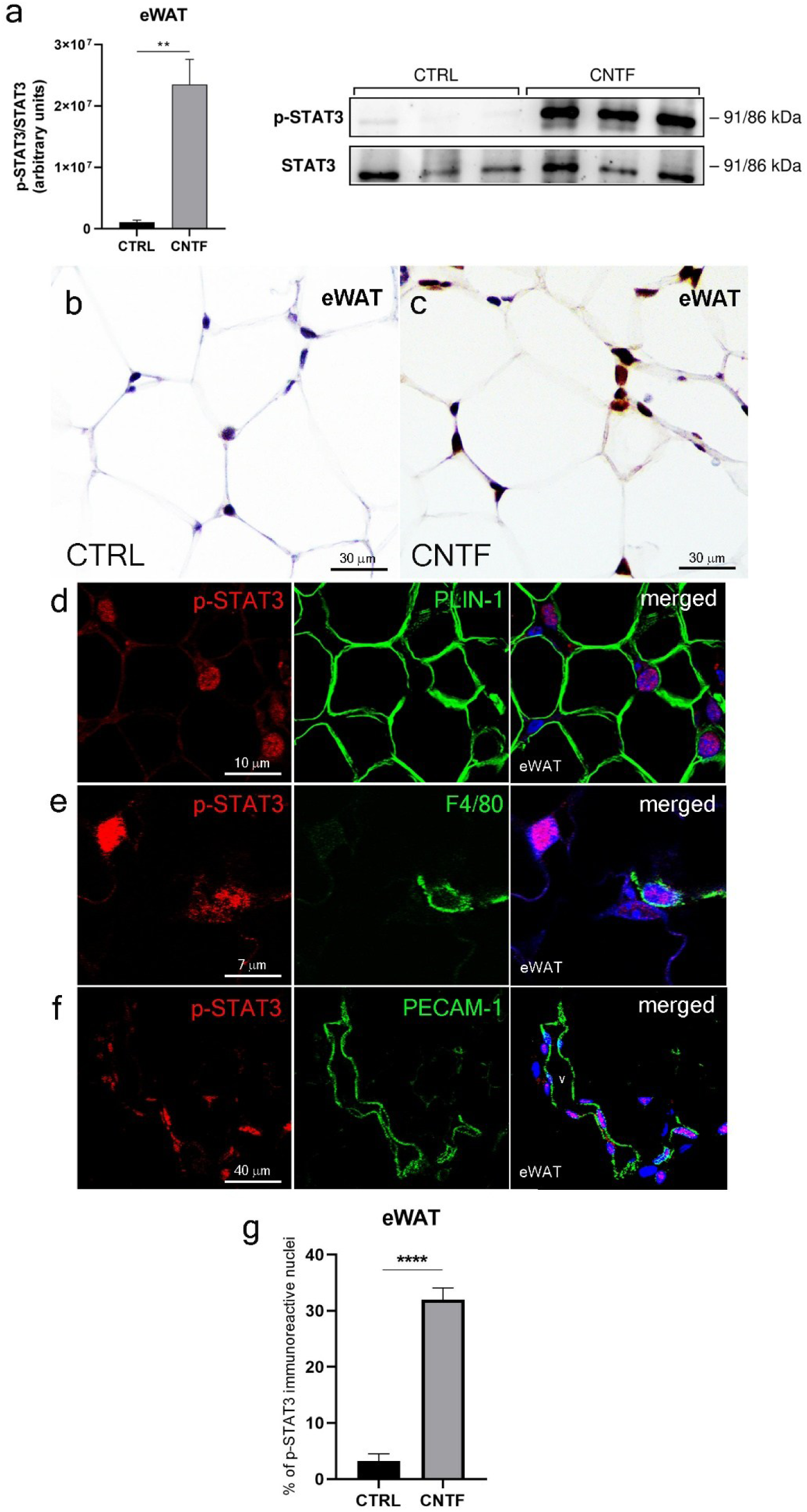
p-STAT3 immunoreactivity after CNTF administration in murine white adipose tissue. a) Representative Western blot and relative densitometric quantification of p-STAT3 protein levels in the epididymal white adipose tissue (eWAT) of saline- (CTRL) and CNTF-treated mice. Total STAT3 was used as a loading control. (b-c) Peroxidase immunohistochemistry for p-STAT3 in the epididymal white adipose tissue after saline or CNTF administration. Double-labelled confocal microscopy for p-STAT3 and the lipid-droplet associated protein (PLIN-1) (d) or the macrophage-marker F4/80 (e) or the endothelial marker Platelet/Endothelial Cell Adhesion Molecule-1 (PECAM-1) (f) in the epididymal white adipose tissue after CNTF administration. (g) Morphometric quantification of p-STAT3-positive nuclei in the eWAT. Data are mean ± SEM (n=3 per group). **p<0.01; ****p < 0.0001; Unpaired t-test.

In small rodents, the interscapular adipose tissue is mainly composed by brown adipocytes, whose thermogenic activity is required to perform thermoregulation, but also affects energy expenditure and body weight (Frontini and Cinti, 2010). Western blot analysis in the iBAT revealed an increased STAT3 phosphorylation following systemic CNTF administration (Fig. 6a). In the iBAT of saline-injected mice, a weak expression of p-STAT3 was detected in the cytoplasm and in very few cellular nuclei by immunohistochemistry (Fig. 6c). A strong STAT3 phosphorylation was observed after CNTF administration in numerous multilocular adipocytes (Fig. 6d). Such response was confirmed by morphometric analysis, revealing a negligible basal activation in saline-treated mice (Fig. 6h and Table 2) compared to a massive response in CNTF-treated mice with the 49 ± 2% of nuclei becoming p-STAT3-positive (vs 0.4 ± 0% in the CTRL, p < 0.0001; Fig. 5h and Table 1). In treated-mice, P-STAT3 specific staining was mainly detected in uncoupling protein 1 (UCP1)-positive brown adipocytes (Fig. 6e). As already observed for the eWAT, however, CNTF administration also led to p-STAT3 immunoreactivity in different cells belonging to the wall of numerous small blood vessels supplying the parenchyma (Fig. 6d, inset).

**Figure 6.**
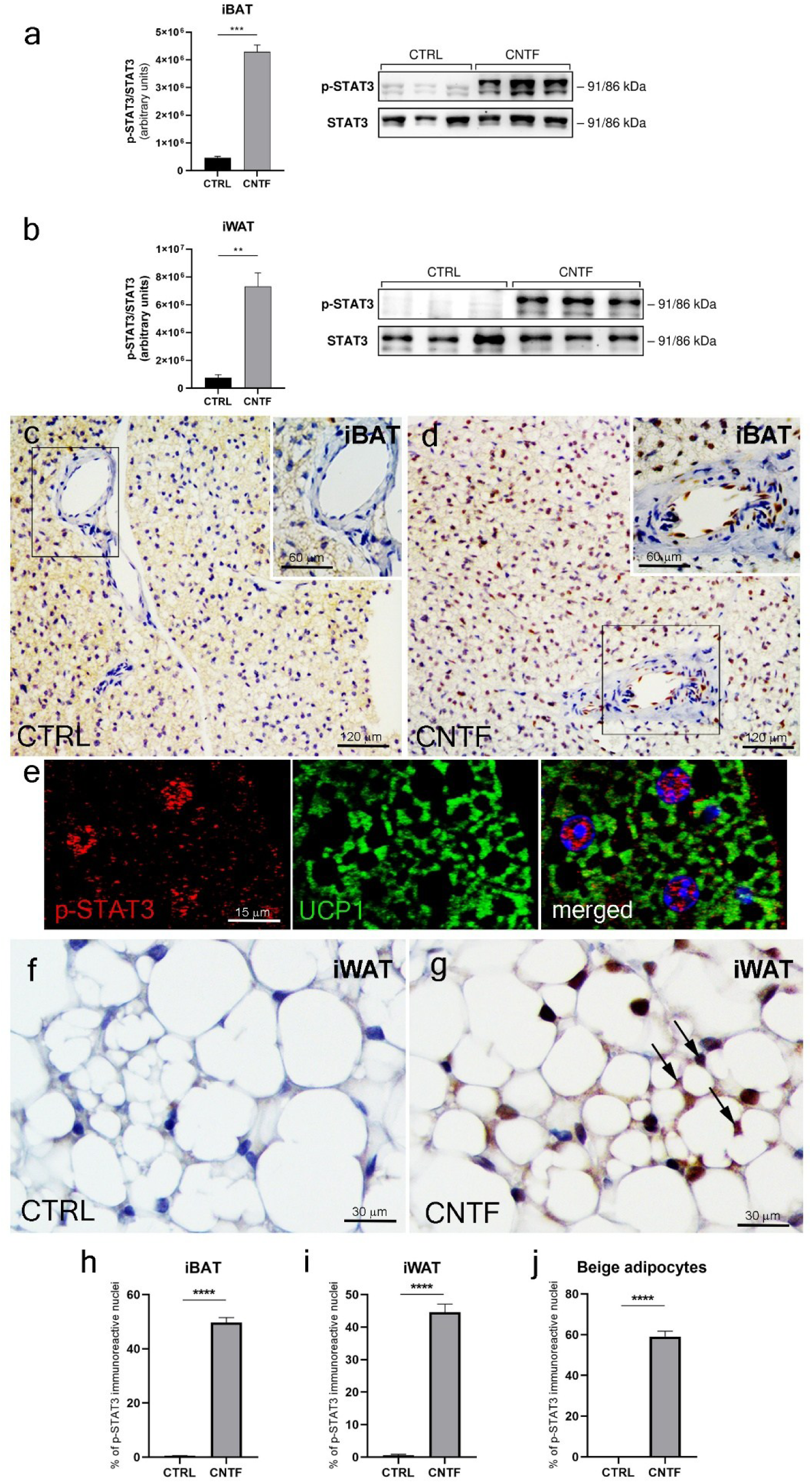
p-STAT3 immunoreactivity after CNTF administration in murine brown and subcutaneous adipose tissue. (a) Western blot and densitometric analysis of p-STAT3 levels in interscapular brown adipose tissue (iBAT) and (b) inguinal white adipose tissue (iWAT) of saline- (CTRL) and CNTF-treated mice. Total STAT3 was used as a loading control. (c-d) Peroxidase immunohistochemistry for p-STAT3 in the brown adipose tissue after saline or CNTF administration. Insets are enlargements of the corresponding framed area in (c) and in (d), with panel (d) showing p-STAT3 immunoreactivity in small blood vessels, particularly arterioles, supplying the parenchyma. (e) Double-labelled confocal microscopy for p-STAT3 and the uncoupling protein 1 (UCP1) after CNTF administration. (f-g) Peroxidase immunohistochemistry for p-STAT3 in the subcutaneous white adipose tissue after saline or CNTF administration. Figure (g) shows a CNTF-dependent STAT3 phosphorylation in paucilocular beige adipocytes (black arrows). (h) Morphometric quantification of the percentage of p-STAT3-positive nuclei in iBAT, iWAT, and beige adipocytes. Data are expressed as mean ± SEM (n=3 per group). **p<0.01; ***p < 0.001, ****p < 0.0001; Unpaired t-test.

In rodents, the subcutaneous iWAT is the largest and most relevant fat depot capable of recruiting beige adipocytes, i.e. brown-like adipocytes, that derive both from *de novo* adipogenesis and/or from the transdifferentiation of white into brown-like cells under proper stimuli (Barbatelli et al., 2010, Wang et al., 2013). Similarly to the iBAT, a strong induction of STAT3 phosphorylation was detected in CNTF-treated mice by Western blotting (Fig. 6b). Consistently, whereas p-STAT3 immunoreactivity was virtually absent in saline-injected mice (Fig 6f), CNTF treatment led to STAT3 phosphorylation in numerous subcutaneous white and beige adipocytes, these latter identified by their distinctive paucilocular morphology (Barbatelli et al., 2010) (Fig. 6g). Morphometric analysis confirmed that p-STAT3-immunoreactive nuclei were nearly absent in the controls (Fig. 6i-j and Table 1), whereas CNTF treatment resulted in a significant increase reaching 44 ± 2% (vs 1 ± 0% in the CTRL, p<0.0001) and 59 ± 2% (vs 0 ± 0% in the CTRL, p<0.0001) of white and beige adipocytes, respectively (Fig. 6i-j and Table 1). As observed for eWAT, p-STAT3 immunoreactivity was detected also in several small blood vessels supplying the tissue, but never in macrophages (data not shown). In conclusion, CNTF administration massively induced STAT3 phosphorylation in visceral white adipocytes of eWAT, in brown adipocytes of iBAT and in white and beige adipocytes of iWAT. In these tissues, CNTF also targeted small blood vessels supplying the adipose depots and, only in the visceral fat, the resident macrophage population.

#### 3.1.5 Skeletal muscle

To assess the effect of CNTF on the skeletal muscle, we evaluated the activation of the JAK/STAT3 pathway in the gastrocnemius muscle. By Western blotting, systemic CNTF administration induced a significant increase in p-STAT3 protein levels compared to saline (Fig. 7a). In the gastrocnemius muscle of the saline-treated mice, p-STAT3 immunoreactivity was virtually absent, with only a few scattered elongated nuclei showing weak basal phosphorylation (Fig. 7b). In contrast, CNTF administration triggered a robust and widespread response (Fig. 7c). Specifically, strong nuclear p-STAT3 immunoreactivity was observed in 36 ± 2% (vs 3 ± 1% in the CTRL, p<0.0001; Fig. 7g and Table 1) of the peripheral nuclei of striated muscle fibers, which are characteristic of these multinucleated syncytial structures. As observed in the adipose tissue, CNTF also induced STAT3 phosphorylation in the endothelial cells of blood vessels supplying the skeletal muscle (data not shown). A particularly striking response was observed in the intramuscular nerves supplying the gastrocnemius, where numerous p-STAT3-positive nuclei were detected in CNTF-treated animals (Fig. 7c, inset).

**Figure 7.**
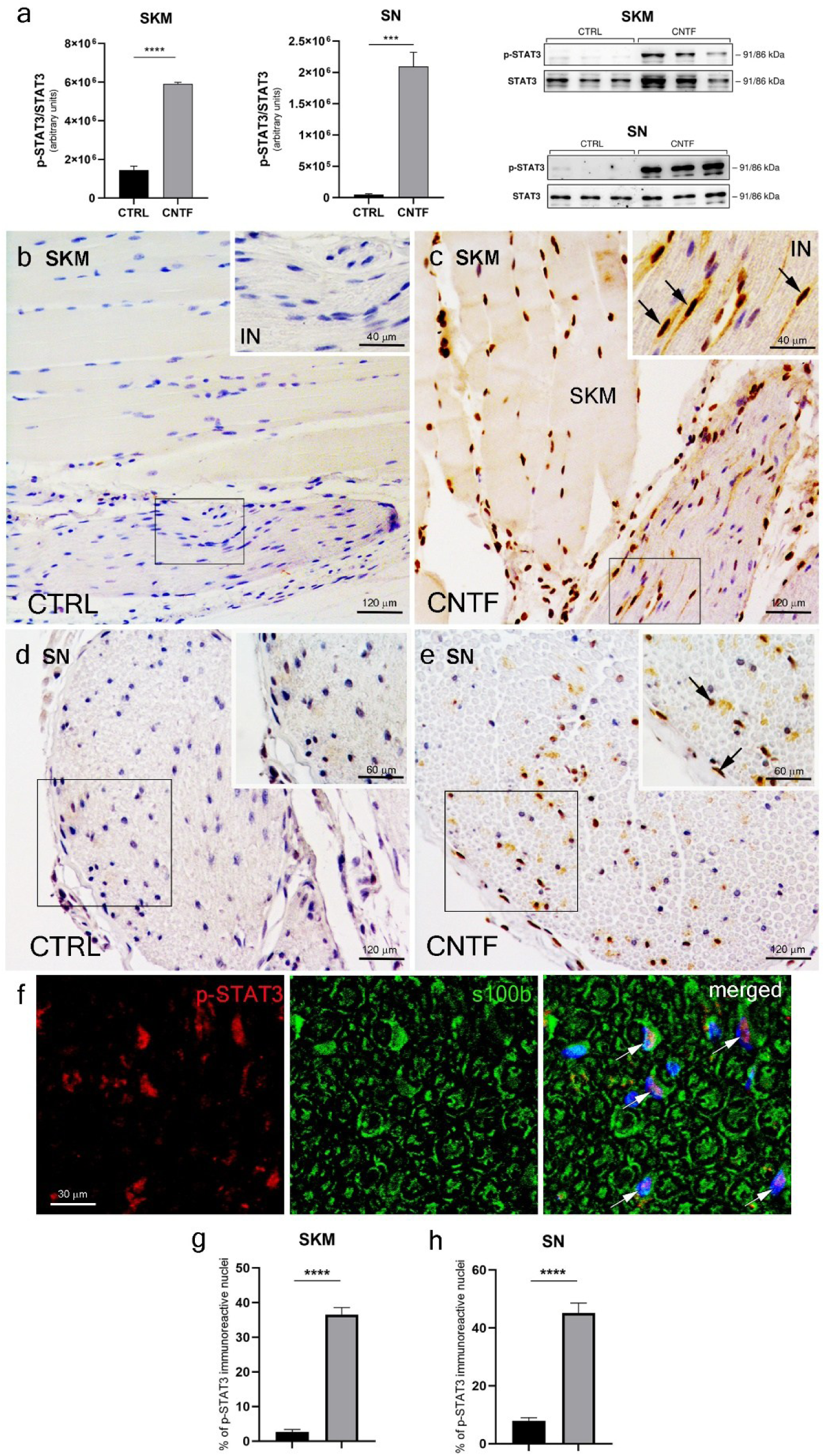
p-STAT3 immunoreactivity after CNTF administration in murine skeletal muscle. (a) Representative Western blot and densitometric quantification of p-STAT3 protein levels in the gastrocnemius skeletal muscle (SKM) and sciatic nerve (SN) of saline- (CTRL) and CNTF-treated mice. Total STAT3 was used as a loading control. (b-c) Peroxidase immunohistochemistry for p-STAT3 in the gastrocnemius after saline or CNTF administration. Insets are enlargements of the corresponding framed area in (b) and (c) and in figure (c) shows p-STAT3 immunoreactivity in peripheral motor nerves innervating the gastrocnemius (black arrows). (d-e) Peroxidase immunohistochemistry for p-STAT3 in the sciatic nerve after saline or CNTF administration. Insets are enlargements of the corresponding framed area in (d) and (e) and in figure (e) shows p-STAT3 immunoreactivity in nerve cells and in cells of the epineurium (black arrows). (f) Double-labelled confocal microscopy for p-STAT3 and the Schwann’s cells marker S100b in sciatic nerve after CNTF administration, showing S100b-positive cells immunoreactive for p-STAT3 (white arrows). (g) Morphometric quantification of the percentage of p-STAT3-immunoreactive nuclei in skeletal muscle and sciatic nerve. Data are expressed as mean ± SEM (n=3 per group). ***p < 0.001, ****p < 0.0001 (Unpaired t-test). IN: intramuscular peripheral nerve.

To further characterize the effect of CNTF on peripheral nerves, we extended our analysis to the sciatic nerve, the prototypical peripheral nerve, containing both sensory and motor axons. Western blot analysis revealed a significant increase of p-STAT3 protein expression in the sciatic nerve of CNTF-treated mice compared to saline (Fig. 7a). Immunohistochemical analysis (Fig. 7d and e) and subsequent morphometric quantification revealed a massive induction of nuclear p-STAT3 in the treated group, reaching 45 ± 3% (vs 8 ± 1% in the CTRL, p<0.0001; Fig. 7h and Table 1). CNTF was found to induce p-STAT3 phosphorylation in the epineurium cells (a layer of connective tissue surrounding nerve fascicles) and in cells colocalizing with S100b (Fig. 7f), a marker for Schwann’s cells, the glial cells involved in axon’s support, survival and myelination. Most CNTF-activated cells located into the intramuscular nerves were S100b-positive Schwann cells (data not shown).

In conclusion, our data demonstrates that systemic CNTF exerts a powerful effect on the skeletal muscle, targeting both the contractile fibers and its vascular component. Importantly, CNTF also acts on skeletal muscle nerve supply, lending support to a possible wide action on peripheral nerves.

## 4. Discussion

CNTF is a neurotrophic factor promoting neuronal survival and differentiation (Arakawa et al., 1990, Oyesiku and Wigston, 1996). Our group has previously demonstrated that exogenous CNTF effectively targets the CNS, bypassing the blood-brain barrier at specific circumventricular organs, such as the median eminence and the area postrema, where it mainly triggers the activation of the JAK2-STAT3 signaling pathway (Severi et al., 2015, Senzacqua et al., 2016, Venema et al., 2020). Furthermore, CNTF was demonstrated to promote leptin entry into hypothalamic feeding centers and to display an anorexigenic activity (Lambert et al., 2001, Sleeman et al., 2003, Venema et al., 2020). At the same time, recent evidence has suggested broader actions for this cytokine outside the CNS, targeting metabolically relevant peripheral organs (Pasquin et al., 2015). It should be noted, however, that much of the current knowledge regarding peripheral CNTF activity stems from *in vitro* studies. In this study, by combining morphological and quantitative approaches, we provided the first comprehensive *in vivo* map of CNTF-responsive cells, using STAT3 phosphorylation as a functional proxy due to the limited availability of reliable antibodies against the Cntfrα (Venema et al., 2020, Zvonic et al., 2003, Severi et al., 2015). Indeed, both central and peripheral effects of exogenous CNTF are significantly blunted or abolished when STAT3 signaling is pharmacologically or genetically inhibited, while remaining largely unaffected by the blockade of alternative CNTF-induced pathways such as ERK1/2-MAPK or AKT (Rezende et al., 2009).

Our *in vivo* mapping revealed a widespread activation of STAT3 signaling across the GI tract. The discrepancy we observed between robust p-STAT3 signaling and negligible *Cntfrα* transcripts in the alimentary system suggests a high signaling efficiency at low receptor densities or, alternatively, the involvement of alternative signaling modes, such as the “trans-signaling” (Rose-John, 2018). This mechanism, shared with other IL-6 family members, allows the circulating complex formed by CNTF and its soluble receptor to activate cells expressing only gp130 and Lifrβ, thereby extending the cytokine’s action even to tissues with low expression of the membrane-bound Cntfrα. Our mapping identified the smooth muscle cells of the muscularis externa as the primary site of STAT3 activation within the GI tract. While previous studies focused on STAT3 in the mucosal epithelium as a mediator of inflammation (Pickert et al., 2009, Bollrath et al., 2009), its activation within the other layers remains largely unexplored *in vivo*. Beyond the muscularis layer, we documented significant p-STAT3 activation in the lamina propria and in the submucosal and myenteric plexuses. These findings align with reports that CNTF promotes the survival of myenteric neurons *in vitro* (Kato et al., 2024). The coordinated activation of STAT3 in the enteric nervous system and contractile layers suggests a direct modulation of the GI motility and provides a novel possible explanation for the nausea and vomiting observed in clinical trials, which were previously attributed solely to central actions in the area postrema (Ettinger et al., 2003, Senzacqua et al., 2016). Our study further extended to the major extramural glands associated with the GI tract, specifically the pancreas and the liver. Regarding the pancreas, our quantitative analysis revealed a prominent p-STAT3 activation in the endocrine pancreas. Our morphological data agree with the present literature reporting that the STAT3 signaling is a well-established transducer of CNTF action in pancreatic islets *in vitro* (Rezende et al., 2007, Rezende et al., 2009). Notably, we showed for the first-time activation of glucagon-producing α-cells alongside insulin-producing β-cells, suggesting coordinated effects on glucose homeostasis involving both cell types. Similarly, in the liver we observed a significant p-STAT3 induction within hepatocytes, Kupffer cells and sinusoids. Since hepatic STAT3 activation is a key regulator of glucose and lipid metabolism—specifically by suppressing gluconeogenic gene expression and enhancing insulin sensitivity (Inoue et al., 2004) — our data added an important interpretative insight into the well-established metabolic effects of the CNTF in the liver (Sleeman et al., 2003, Cui et al., 2017). A notable finding of our mapping is the recruitment of STAT3 signaling within several immune populations, including adipose tissue macrophages. As described by Akira et al. (Akira, 2000), STAT3 plays a key role in immune cells, often mediating anti-inflammatory and pro-survival roles. In the context of adipose tissue, STAT3 activation occurred in both macrophages and adipocytes, where it has been linked to lipolysis and browning (Cernkovich et al., 2008, Derecka et al., 2012), suggesting that the well-described anti-obesity effects of CNTF *in vitro* (Zvonic et al., 2003, Perugini et al., 2019) may involve a direct modulation of the adipose tissue microenvironment and its inflammatory state *in vivo*. For the first time, we documented a significant JAK2-STAT3 activation in beige adipocytes, further suggesting that CNTF may involve the recruitment of thermogenic programs across different adipose depots. Lastly, as a major site of energy substrate utilization, we showed a significant STAT3 phosphorylation in the skeletal muscle, where CNTF is known to improve insulin sensitivity, increase protein synthesis (Wang and Forsberg, 2000), and enhance fatty acid utilization (Zvonic et al., 2003, Watt et al., 2006). Our study showed that, within skeletal muscle, STAT3 signaling is also activated in Schwann cells of peripheral nerves supplying the tissue, including the sciatic nerve. Such response in glial cells—a major source of endogenous CNTF—suggests a potential autocrine/paracrine loop (Lee et al., 1995).

The biological outcome of STAT3 signaling is critically dependent on the dosage and its temporal dynamics. In our model, the acute recruitment of STAT3 signaling reflects a physiological stimulus generally considered homeostatic and protective, differing significantly from the persistent overactivation associated with chronic inflammatory diseases and metabolic dysfunction (Haikarainen et al., 2026, Rose-John, 2018).

While our study defines an anatomical fingerprint of acute CNTF administration, we acknowledge some limitations. First, as our study is observational in nature, the functional implications of these findings are inferred based on existing literature. Second, our analysis was restricted to male subjects, thus not accounting for sex-specific responses previously documented (Perugini et al., 2022, Colleluori et al., 2025). Lastly, an additional limitation is that CNTF can induce IL-6 release, which in turn activates the same signaling cascade. Although this process occurs later than the 45 minute window used in our protocol, we cannot entirely exclude the influence of the endogenous cytokine background on STAT3 phosphorylation (Hu et al., 2020). In conclusion, this anatomical mapping underscores the necessity of considering cell-type-specific responses to understand the therapeutic potential and side-effect profile of CNTF-based treatments. By identifying novel peripheral targets, we provide a morphological framework for future functional studies on cytokine-mediated metabolic regulation.

## 5. ACKNOWLEDGMENTS

This work was supported by grants from Fondazione Cariverona (Bando Progetti Scientifici di Eccellenza 2018) to AG, Fondazione di Medicina Molecolare e Terapia Cellulare to AG, ESPEN Research Fellowship Grant and EFSD Rising Star Grant to GC, and Marche Polytechnic University (Contributi Ricerca Scientifica di Ateneo).

## 6. AUTHOR CONTRIBUTIONS

Chiara Galli: concept/design, acquisition of data, data analysis/interpretation, drafting of the manuscript; Georgia Colleluori: acquisition of data, data analysis/interpretation, drafting of the manuscript; Jessica Perugini, Edoardo Scopini, Ilenia Severi and Gaia Grandin: acquisition of data, data analysis/interpretation; Antonio Giordano: concept/design, critical revision of the manuscript and approval of the article.

## REFERENCES

Aaronson, D. S. & Horvath, C. M. 2002. A road map for those who don’t know JAK-STAT. Science, 296, 1653–5.

Adler, R., Landa, K. B., Manthorpe, M. & Varon, S. 1979. Cholinergic neuronotrophic factors: intraocular distribution of trophic activity for ciliary neurons. Science, 204, 1434–6.

Akira, S. 2000. Roles of STAT3 defined by tissue-specific gene targeting. Oncogene, 19, 2607–11.

Al-Adsani, A. M., Barhoush, S. A., Bastaki, N. K., Al-Bustan, S. A. & Al-Qattan, K. K. 2022. Comparing and Optimizing RNA Extraction from the Pancreas of Diabetic and Healthy Rats for Gene Expression Analyses. Genes (Basel), 13.

Anderson, K. D., Lambert, P. D., Corcoran, T. L., Murray, J. D., Thabet, K. E., Yancopoulos, G. D. & Wiegand, S. J. 2003. Activation of the hypothalamic arcuate nucleus predicts the anorectic actions of ciliary neurotrophic factor and leptin in intact and gold thioglucose-lesioned mice. J Neuroendocrinol, 15, 649–60.

Arakawa, Y., Sendtner, M. & Thoenen, H. 1990. Survival effect of ciliary neurotrophic factor (CNTF) on chick embryonic motoneurons in culture: comparison with other neurotrophic factors and cytokines. J Neurosci, 10, 3507–15.

Augereau, C., Lemaigre, F. P. & Jacquemin, P. 2016. Extraction of high-quality RNA from pancreatic tissues for gene expression studies. Anal Biochem, 500, 60–2.

Azevedo-Pouly, A. C., Elgamal, O. A. & Schmittgen, T. D. 2014. RNA isolation from mouse pancreas: a ribonuclease-rich tissue. J Vis Exp, e51779.

Barbatelli, G., Murano, I., Madsen, L., Hao, Q., Jimenez, M., Kristiansen, K., Giacobino, J. P., DE Matteis, R. & Cinti, S. 2010. The emergence of cold-induced brown adipocytes in mouse white fat depots is determined predominantly by white to brown adipocyte transdifferentiation. Am J Physiol Endocrinol Metab, 298, E1244–53.

Bluher, S., Moschos, S., Bullen, J., JR., Kokkotou, E., Maratos-Flier, E., Wiegand, S. J., Sleeman, M. W. & Mantzoros, C. S. 2004. Ciliary neurotrophic factorAx15 alters energy homeostasis, decreases body weight, and improves metabolic control in diet-induced obese and UCP1-DTA mice. Diabetes, 53, 2787–96.

Bollrath, J., Phesse, T. J., Von Burstin, V. A., Putoczki, T., Bennecke, M., Bateman, T., Nebelsiek, T., Lundgren-May, T., Canli, O., Schwitalla, S., Matthews, V., Schmid, R. M., Kirchner, T., Arkan, M. C., Ernst, M. & Greten, F. R. 2009. gp130-mediated Stat3 activation in enterocytes regulates cell survival and cell-cycle progression during colitis-associated tumorigenesis. Cancer Cell, 15, 91–102.

Cernkovich, E. R., Deng, J., Bond, M. C., Combs, T. P. & Harp, J. B. 2008. Adipose-specific disruption of signal transducer and activator of transcription 3 increases body weight and adiposity. Endocrinology, 149, 1581–90.

Colleluori, G., Viola, V., Bathina, S., Armamento-Villareal, R., Qualls, C., Giordano, A. & Villareal, D. T. 2025. Effect of aerobic or resistance exercise, or both on insulin secretion, ciliary neurotrophic factor, and insulin-like growth factor-1 in dieting older adults with obesity. Clin Nutr, 51, 50–62.

Cui, M. X., Yang, L. N., Wang, X. X., Wang, L., Li, R. L., Han, W. & Wu, Y. J. 2017. Alleviative effect of ciliary neurotrophic factor analogue on high fat-induced hepatic steatosis is partially independent of the central regulation. Clin Exp Pharmacol Physiol, 44, 395–402.

Davis, S., Aldrich, T. H., Stahl, N., Pan, L., Taga, T., Kishimoto, T., Ip, N. Y. & Yancopoulos, G. D. 1993. LIFR beta and gp130 as heterodimerizing signal transducers of the tripartite CNTF receptor. Science, 260, 1805–8.

Derecka, M., Gornicka, A., Koralov, S. B., Szczepanek, K., Morgan, M., Raje, V., Sisler, J., Zhang, Q., Otero, D., Cichy, J., Rajewsky, K., Shimoda, K., Poli, V., Strobl, B., Pellegrini, S., Harris, T. E., Seale, P., Russell, A. P., Mcainch, A. J., O’brien, P. E., Keller, S. R., Croniger, C. M., Kordula, T. & Larner, A. C. 2012. Tyk2 and Stat3 regulate brown adipose tissue differentiation and obesity. Cell Metab, 16, 814–24.

Dittrich, F., Thoenen, H. & Sendtner, M. 1994. Ciliary neurotrophic factor: pharmacokinetics and acute-phase response in rat. Ann Neurol, 35, 151–63.

Ettinger, M. P., Littlejohn, T. W., Schwartz, S. L., Weiss, S. R., Mcilwain, H. H., Heymsfield, S. B., Bray, G. A., Roberts, W. G., Heyman, E. R., Stambler, N., Heshka, S., Vicary, C. & Guler, H. P. 2003. Recombinant variant of ciliary neurotrophic factor for weight loss in obese adults: a randomized, dose-ranging study. Jama, 289, 1826–32.

Frontini, A., Bertolotti, P., Tonello, C., Valerio, A., Nisoli, E., Cinti, S. & Giordano, A. 2008. Leptin-dependent STAT3 phosphorylation in postnatal mouse hypothalamus. Brain Res, 1215, 105–15.

Frontini, A. & Cinti, S. 2010. Distribution and development of brown adipocytes in the murine and human adipose organ. Cell Metab, 11, 253–6.

Ghanemi, A. & St-Amand, J. 2018. Interleukin-6 as a “metabolic hormone”. Cytokine, 112, 132–136.

Haikarainen, T., Virtanen, A. T., Cravatt, B. F. & Silvennoinen, O. 2026. Pharmacological targeting of the JAK-STAT pathway: new concepts and emerging indications. Nat Rev Drug Discov, 25, 268–289.

Heinrich, P. C., Behrmann, I., Muller-Newen, G., Schaper, F. & Graeve, L. 1998. Interleukin-6-type cytokine signalling through the gp130/Jak/STAT pathway. Biochem J, 334 ( Pt 2), 297–314.

Hirasawa, A., Tsumaya, K., Awaji, T., Katsuma, S., Adachi, T., Yamada, M., Sugimoto, Y., Miyazaki, S. & Tsujimoto, G. 2005. Free fatty acids regulate gut incretin glucagon-like peptide-1 secretion through GPR120. Nat Med, 11, 90–4.

Holst, J. J. 2004. On the physiology of GIP and GLP-1. Horm Metab Res, 36, 747–54.

Hu, Z., Deng, N., Liu, K., Zhou, N., Sun, Y. & Zeng, W. 2020. CNTF-STAT3-IL-6 Axis Mediates Neuroinflammatory Cascade across Schwann Cell-Neuron-Microglia. Cell Rep, 31, 107657.

Hubschle, T., Thom, E., Watson, A., Roth, J., Klaus, S. & Meyerhof, W. 2001. Leptin-induced nuclear translocation of STAT3 immunoreactivity in hypothalamic nuclei involved in body weight regulation. J Neurosci, 21, 2413–24.

Inoue, H., Ogawa, W., Ozaki, M., Haga, S., Matsumoto, M., Furukawa, K., Hashimoto, N., Kido, Y., Mori, T., Sakaue, H., Teshigawara, K., Jin, S., Iguchi, H., Hiramatsu, R., Leroith, D., Takeda, K., Akira, S. & Kasuga, M. 2004. Role of STAT-3 in regulation of hepatic gluconeogenic genes and carbohydrate metabolism in vivo. Nat Med, 10, 168–74.

Ip, N. Y., Mcclain, J., Barrezueta, N. X., Aldrich, T. H., Pan, L., Li, Y., Wiegand, S. J., Friedman, B., Davis, S. & Yancopoulos, G. D. 1993. The alpha component of the CNTF receptor is required for signaling and defines potential CNTF targets in the adult and during development. Neuron, 10, 89–102.

Judd, L. M., Bredin, K., Kalantzis, A., Jenkins, B. J., Ernst, M. & Giraud, A. S. 2006. STAT3 activation regulates growth, inflammation, and vascularization in a mouse model of gastric tumorigenesis. Gastroenterology, 131, 1073–85.

Kato, R., Yamamoto, T., Ogata, H., Miyata, K., Hayashi, S., Gershon, M. D. & Kadowaki, M. 2024. Indigenous gut microbiota constitutively drive release of ciliary neurotrophic factor from mucosal enteric glia to maintain the homeostasis of enteric neural circuits. Front Immunol, 15, 1372670.

Kolodziejczyk, A. A. & Lonnberg, T. 2018. Global and targeted approaches to single-cell transcriptome characterization. Brief Funct Genomics, 17, 209–219.

Lambert, P. D., Anderson, K. D., Sleeman, M. W., Wong, V., Tan, J., Hijarunguru, A., Corcoran, T. L., Murray, J. D., Thabet, K. E., Yancopoulos, G. D. & Wiegand, S. J. 2001. Ciliary neurotrophic factor activates leptin-like pathways and reduces body fat, without cachexia or rebound weight gain, even in leptin-resistant obesity. Proc Natl Acad Sci U S A, 98, 4652–7.

Lee, D. A., Zurawel, R. H. & Windebank, A. J. 1995. Ciliary neurotrophic factor expression in Schwann cells is induced by axonal contact. J Neurochem, 65, 564–8.

Lertkiatmongkol, P., Liao, D., Mei, H., Hu, Y. & Newman, P. J. 2016. Endothelial functions of platelet/endothelial cell adhesion molecule-1 (CD31). Curr Opin Hematol, 23, 253–9.

Liu, Y., Liu, H., Meyer, C., Li, J., Nadalin, S., Konigsrainer, A., Weng, H., Dooley, S. & TEN Dijke, P. 2013. Transforming growth factor-beta (TGF-beta)-mediated connective tissue growth factor (CTGF) expression in hepatic stellate cells requires Stat3 signaling activation. J Biol Chem, 288, 30708–30719.

Lumeng, C. N., Bodzin, J. L. & Saltiel, A. R. 2007. Obesity induces a phenotypic switch in adipose tissue macrophage polarization. J Clin Invest, 117, 175–84.

Miller, R. G., Petajan, J. H., Bryan, W. W., Armon, C., Barohn, R. J., Goodpasture, J. C., Hoagland, R. J., Parry, G. J., Ross, M. A. & Stromatt, S. C. 1996. A placebo-controlled trial of recombinant human ciliary neurotrophic (rhCNTF) factor in amyotrophic lateral sclerosis. rhCNTF ALS Study Group. Ann Neurol, 39, 256–60.

Miyaoka, Y., Tanaka, M., Naiki, T. & Miyajima, A. 2006. Oncostatin M inhibits adipogenesis through the RAS/ERK and STAT5 signaling pathways. J Biol Chem, 281, 37913–20.

Murakami, M., Kamimura, D. & Hirano, T. 2019. Pleiotropy and Specificity: Insights from the Interleukin 6 Family of Cytokines. Immunity, 50, 812–831.

Naito, M., Hasegawa, G., Ebe, Y. & Yamamoto, T. 2004. Differentiation and function of Kupffer cells. Med Electron Microsc, 37, 16–28.

Oyesiku, N. M. & Wigston, D. J. 1996. Ciliary neurotrophic factor stimulates neurite outgrowth from spinal cord neurons. J Comp Neurol, 364, 68–77.

Pasquin, S., Sharma, M. & Gauchat, J. F. 2015. Ciliary neurotrophic factor (CNTF): New facets of an old molecule for treating neurodegenerative and metabolic syndrome pathologies. Cytokine Growth Factor Rev, 26, 507–15.

Perugini, J., Di Mercurio, E., Giuliani, A., Sabbatinelli, J., Bonfigli, A. R., Tortato, E., Severi, I., Cinti, S., Olivieri, F., LE Roux, C. W., Gesuita, R. & Giordano, A. 2022. Ciliary neurotrophic factor is increased in the plasma of patients with obesity and its levels correlate with diabetes and inflammation indices. Sci Rep, 12, 8331.

Perugini, J., DI Mercurio, E., Tossetta, G., Severi, I., Monaco, F., Reguzzoni, M., Tomasetti, M., Dani, C., Cinti, S. & Giordano, A. 2019. Biological Effects of Ciliary Neurotrophic Factor on hMADS Adipocytes. Front Endocrinol (Lausanne), 10, 768.

Pickert, G., Neufert, C., Leppkes, M., Zheng, Y., Wittkopf, N., Warntjen, M., Lehr, H. A., Hirth, S., Weigmann, B., Wirtz, S., Ouyang, W., Neurath, M. F. & Becker, C. 2009. STAT3 links IL-22 signaling in intestinal epithelial cells to mucosal wound healing. J Exp Med, 206, 1465–72.

Rezende, L. F., Stoppiglia, L. F., Souza, K. L., Negro, A., Langone, F. & Boschero, A. C. 2007. Ciliary neurotrophic factor promotes survival of neonatal rat islets via the BCL-2 anti-apoptotic pathway. J Endocrinol, 195, 157–65.

Rezende, L. F., Vieira, A. S., Negro, A., Langone, F. & Boschero, A. C. 2009. Ciliary neurotrophic factor (CNTF) signals through STAT3-SOCS3 pathway and protects rat pancreatic islets from cytokine-induced apoptosis. Cytokine, 46, 65–71.

Rose-John, S. 2018. Interleukin-6 Family Cytokines. Cold Spring Harb Perspect Biol, 10.

Sendtner, M., Carroll, P., Holtmann, B., Hughes, R. A. & Thoenen, H. 1994. Ciliary neurotrophic factor. J Neurobiol, 25, 1436–53.

Senzacqua, M., Severi, I., Perugini, J., Acciarini, S., Cinti, S. & Giordano, A. 2016. Action of Administered Ciliary Neurotrophic Factor on the Mouse Dorsal Vagal Complex. Front Neurosci, 10, 289.

Severi, I., Senzacqua, M., Mondini, E., Fazioli, F., Cinti, S. & Giordano, A. 2015. Activation of transcription factors STAT1 and STAT5 in the mouse median eminence after systemic ciliary neurotrophic factor administration. Brain Res, 1622, 217–29.

Simi, A. & Ibanez, C. F. 2010. Assembly and activation of neurotrophic factor receptor complexes. Dev Neurobiol, 70, 323–31.

Sleeman, M. W., Anderson, K. D., Lambert, P. D., Yancopoulos, G. D. & Wiegand, S. J. 2000. The ciliary neurotrophic factor and its receptor, CNTFR alpha. Pharm Acta Helv, 74, 265–72.

Sleeman, M. W., Garcia, K., Liu, R., Murray, J. D., Malinova, L., Moncrieffe, M., Yancopoulos, G. D. & Wiegand, S. J. 2003. Ciliary neurotrophic factor improves diabetic parameters and hepatic steatosis and increases basal metabolic rate in db/db mice. Proc Natl Acad Sci U S A, 100, 14297–302.

Steiner, D. J., Kim, A., Miller, K. & Hara, M. 2010. Pancreatic islet plasticity: interspecies comparison of islet architecture and composition. Islets, 2, 135–45.

Tabula Muris, C. 2018. Single-cell transcriptomics of 20 mouse organs creates a Tabula Muris. Nature, 562, 367–372.

Venema, W., Severi, I., Perugini, J., DI Mercurio, E., Mainardi, M., Maffei, M., Cinti, S. & Giordano, A. 2020. Ciliary Neurotrophic Factor Acts on Distinctive Hypothalamic Arcuate Neurons and Promotes Leptin Entry Into and Action on the Mouse Hypothalamus. Front Cell Neurosci, 14, 140.

Wang, M. C. & Forsberg, N. E. 2000. Effects of ciliary neurotrophic factor (CNTF) on protein turnover in cultured muscle cells. Cytokine, 12, 41–8.

Wang, Q. A., Tao, C., Gupta, R. K. & Scherer, P. E. 2013. Tracking adipogenesis during white adipose tissue development, expansion and regeneration. Nat Med, 19, 1338–44.

Watt, M. J., Dzamko, N., Thomas, W. G., Rose-John, S., Ernst, M., Carling, D., Kemp, B. E., Febbraio, M. A. & Steinberg, G. R. 2006. CNTF reverses obesity-induced insulin resistance by activating skeletal muscle AMPK. Nat Med, 12, 541–8.

Yamada, T., Oka, Y. & Katagiri, H. 2008. Inter-organ metabolic communication involved in energy homeostasis: potential therapeutic targets for obesity and metabolic syndrome. Pharmacol Ther, 117, 188–98.

Zvonic, S., Cornelius, P., Stewart, W. C., Mynatt, R. L. & Stephens, J. M. 2003. The regulation and activation of ciliary neurotrophic factor signaling proteins in adipocytes. J Biol Chem, 278, 2228–35.

