## Supplementary tables and figures for "The ciliary neurotrophic factor induces Stat3 phosphorylation in distinctive cytotypes of organs involved in body metabolism: an immunohistochemical study"

### SUPPLEMENTARY MATERIALS

**Table S1**. Taqman probes

| Target Gene | Assay ID |
|---|---|
| *Cntfrα* | Mm00516693_m1 |
| *Lifrβ* | Mm00442942_m1 |
| *Gp130* | Mm00439665_m1 |
| *Tbp* | Mm00446973_m1 |

**Table S2a:** Primary antibodies

| Antibody | Source | Dilution | Brand | Application |
|---|---|---|---|---|
| p-STAT3 (Tyr 705) | Monoclonal rabbit | 1:600 | Cell Signalling (9145), RRID: AB_2491009 | IHC (All); IF (mesenteric gu; pancreas; sciatic nerve) |
| p-STAT3 (Tyr 705) | Monoclonal mouse | 1:200 | Cell Signalling (4113), RRID: AB_2198588 | IF (mesenteric gut; liver, white and brown adipose tissues); WB |
| STAT3 | Polyclonal rabbit | 1:1000 | Cell Signalling (9132), RRID:AB_331588 | WB |
| PGP9.5 | Polyclonal rabbit | 1:200 | Invitrogen (38-1000), RRID: AB_2533355 | IF |
| F4/80 | Monoclonal rabbit | 1:300 | Cell Signalling (70076), RRID: AB_2799771 | IF |
| PLIN-1 | Polyclonal rabbit | 1:500 | Abcam (ab3526), RRID: AB_2167274 | IF |
| CD31 (PECAM-1) | Monoclonal rabbit | 1:50 | Cell Signalling (77699), RRID: AB_2722705 | IF |
| UCP1 | Polyclonal rabbit | 1:800 | Merck (662045) | IF |
| Insulin | Polyclonal guinea pig | 1:400 | Invitrogen (PA1-26938), RRID: AB_794668 | IF |
| Glucagon | Monoclonal mouse | 1:400 | Santa Cruz (sc-57171), RRID: AB_783558 | IF |

| | | | | |
|---|---|---|---|---|
| S100b | Polyclonal guinea pig | 1:200 | Synaptic System (287 004), RRID: AB_2620025 | IF |
| GLP-1 | Polyclonal guinea pig | 1:500 | Synaptic System (471 005), RRID: AB_2924957 | IF |
| GIP | Polyclonal rabbit | 1:500 | Synaptic System (514003), RRID: AB_3662043 | IF |

IHC=peroxidase immunohistochemistry; IF=immunofluorescence; WB= western blot

**Table S2b:** Secondary antibodies

| **Conjugate** | **Source** | **Target** | **Dilution** | **Brand** | **Application** |
|---|---|---|---|---|---|
| HRP | Goat | Mouse | 1:5000 | Bio-Rad (1706516), RRID: AB_11125547 | WB |
| HRP | Goat | Rabbit | 1:5000 | Bio-Rad (1706515), RRID: AB_11125142 | WB |
| Anti-rabbit IgG biotinylated | Goat | Anti-rabbit | 1:200 | Vector Laboratories (BA-1000), RRID: AB_2313606 | IHC |
| Alexa Fluor Cy3 | Donkey | Anti-mouse | 1:200 | Jackson ImmunoResearch (715-166-151), RRID: AB_2340817 | IF |
| Alexa Fluor 488 | Donkey | Anti-rabbit | 1:400 | Invitrogen (A-21206), RRID: AB_2535792 | IF |
| Alexa Fluor 488 | Donkey | Anti-guinea pig | 1:200 | Jackson ImmunoResearch (706-546-148), | IF |

| | | | | | |
|---|---|---|---|---|---|
| | | | | RRID: AB_2340473 | |
| Alexa Fluor 647 | Donkey | Anti-mouse | 1:200 | Jackson ImmunoResearch (706-605-148), RRID: AB_2340476 | IF |

IHC=peroxidase immunohistochemistry; IF=immunofluorescence; WB= western blot

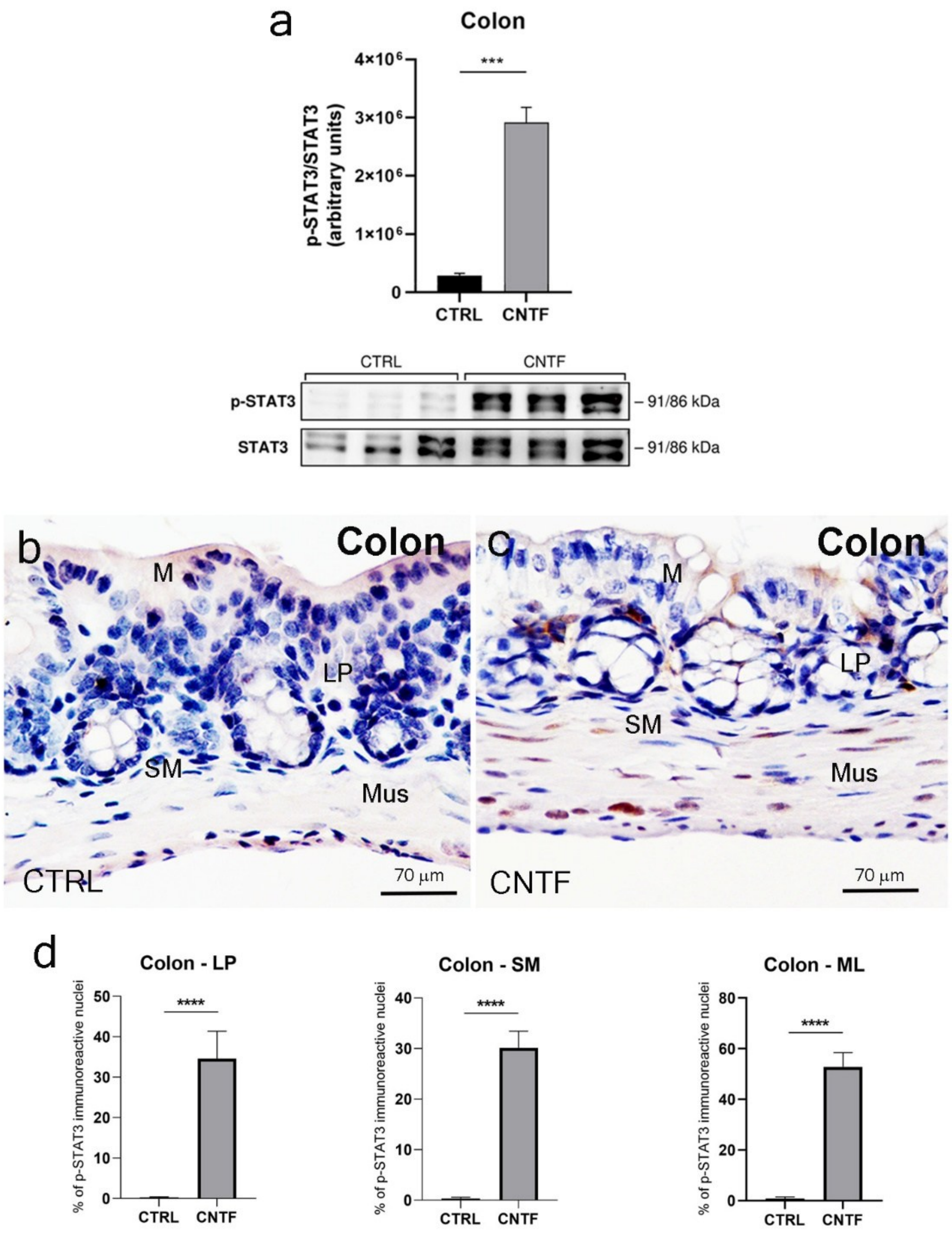


***Figure S1. p-STAT3 immunoreactivity after CNTF administration in murine colon.*** (a) Western blot and densitometric analysis of p-STAT3 protein levels in the colon of saline- (CTRL) and CNTF-treated mice. Total STAT3 was used as a loading control. (b-c) Immunohistochemical staining showing the distribution of p-STAT3-positive nuclei in the colon. (d) Morphometric analysis of p-STAT3-positive nuclei across the lamina propria (LP), submucosa (SM), and muscularis layer (ML) of the colon. Data are mean ± SEM (n=3 animals per group). ***$p < 0.001$, ****$p < 0.0001$ (Unpaired t-test).

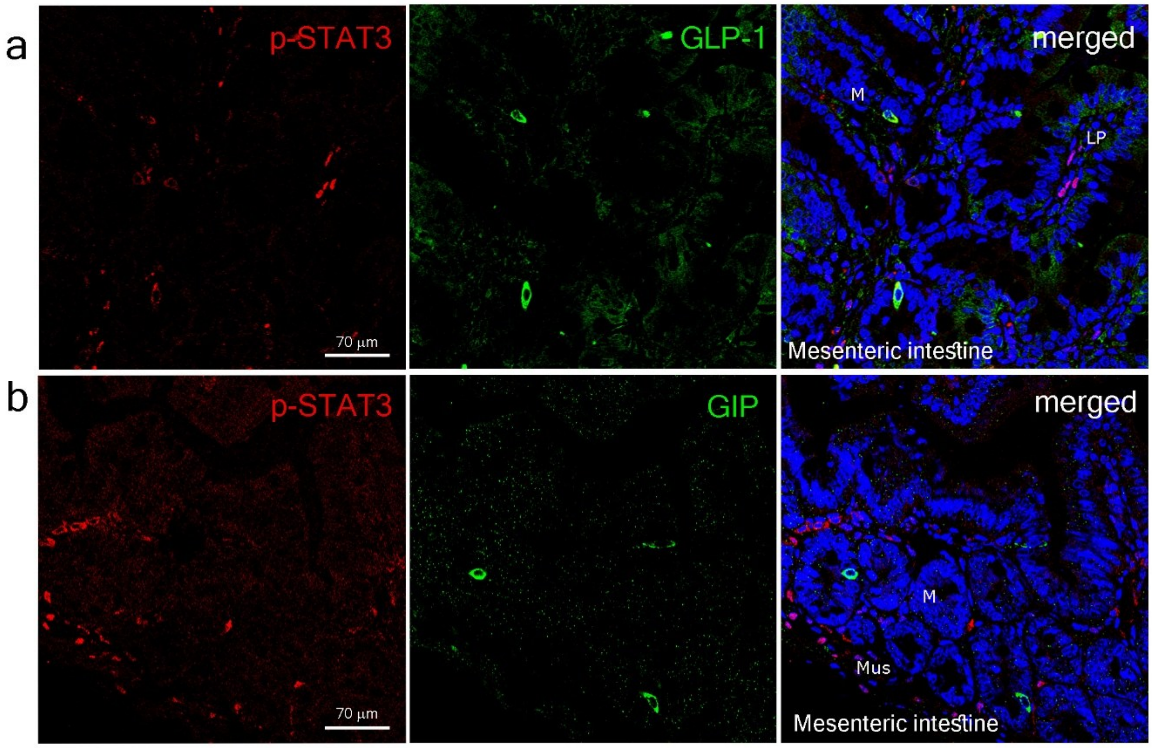


***Figure S2. CNTF response in incretin-producing enteroendocrine cells in murine intestine.*** Double-labelled confocal microscopy for p-STAT3 and the glucagon-like peptide 1 (GLP-1) (a), or the gastric inhibitory peptide (GIP) (b) in the mesenteric gut after CNTF administration. M=mucosa; LP=lamina propria; Mus=muscularis externa.
